# Inferring Adaptive Introgression Using Hidden Markov Models

**DOI:** 10.1101/2020.08.02.232934

**Authors:** Jesper Svedberg, Vladimir Shchur, Solomon Reinman, Rasmus Nielsen, Russell Corbett-Detig

## Abstract

Adaptive introgression - the flow of adaptive genetic variation between species or populations - has attracted significant interest in recent years and it has been implicated in a number of cases of adaptation, from pesticide resistance and immunity, to local adaptation. Despite this, methods for identification of adaptive introgression from population genomic data are lacking. Here, we present Ancestry_HMM-S, a Hidden Markov Model based method for identifying genes undergoing adaptive introgression and quantifying the strength of selection acting on them. Through extensive validation, we show that this method performs well on moderately sized datasets for realistic population and selection parameters. We apply Ancestry_HMM-S to a dataset of an admixed *Drosophila melanogaster* population from South Africa and we identify 17 loci which show signatures of adaptive introgression, four of which have previously been shown to confer resistance to insecticides. Ancestry_HMM-S provides a powerful method for inferring adaptive introgression in datasets that are typically collected when studying admixed populations. This method will enable powerful insights into the genetic consequences of admixture across diverse populations. Ancestry_HMM-S can be downloaded from https://github.com/jesvedberg/Ancestry_HMM-S/.

## Introduction

It is becoming increasingly clear that admixture, gene flow between genetically divergent populations, is a common phenomenon in nature. In some cases, introgressed genetic material confers a selective advantage for individuals in the recipient population, commonly referred to as adaptive introgression, and it is thought to underlie the evolution of numerous adaptive phenotypes (Hedrick 2013; Racimo et al. 2015; Suarez-Gonzalez, Adriana et al. 2018), for example pesticide resistance in mice (Song et al. 2011) and mosquitos (Norris et al. 2015), and complex mimicry patterns in *Heliconius* butterflies (The Heliconius Genome Consortium, 2012). Perhaps the most famous example is the introgression of an allele of *EPAS1* from archaic Denisovans into a modern human population, where the Denisovan allele is thought to have increased in frequency in Tibet due to higher fitness at high altitudes (Huerta-Sánchez et al. 2014; Jeong et al. 2014; Racimo et al. 2015). Admixture therefore has the potential to facilitate adaptive phenotypic outcomes across diverse populations and is rapidly emerging as one of the fundamental drivers of natural selection (Hedrick 2013; Suarez-Gonzalez, Adriana et al. 2018).

Recent admixture is thought to be an important evolutionary force in *D. melanogaster* as well. Populations of this species migrated out from sub-Saharan Africa to colonize the rest of the world approximately 10,000-15,000 years ago (Thornton and Andolfatto 2006). During this expansion, the population that left Africa experienced a dramatic bottleneck that reshaped haplotypic variation across the genome, resulting in decreased diversity and extended linkage disequilibrium (Thornton and Andolfatto 2006; Pool et al. 2012). More recently, descendants of the ancestral and derived populations have admixed in several locations across the world, and these have been the subjects of numerous previous analyses of admixture and local ancestry (Pool et al. 2012; Kao et al. 2015; Lack et al. 2015; Pool 2015; Bergland et al. 2016; Lack et al. 2016; Corbett-Detig and Nielsen 2017; Medina et al. 2018). In particular, the population history for one large admixed sample from South Africa (Lack et al. 2015) is consistent with a simple admixture model where cosmopolitan ancestry introgressed into this population once approximately 30 years prior to sampling (Medina et al. 2018).

Although there have been numerous investigations into the factors that cause nascent reproductive isolation between populations (*e.g*. Coyne and Orr, 2004) and genome-wide signatures of selection (Kolaczkowski et al. 2011; Langley et al. 2012; Reinhardt et al. 2014; Garud et al. 2015), comparatively little work has focused on adaptive outcomes resulting from admixture in *D. melanogaster*. As in other species, pesticides are a major driver of selection, and resistance factors can quickly spread in populations when pesticides are introduced, either from standing genetic variation or from *de novo* mutations (Karasov et al. 2010; Garud et al. 2015). In *D. melanogaster*, specific alleles of several different genes are known to confer resistance to common pesticides. For instance, alleles of several *Cyp6* Cytochrome P450 genes are implicated in DDT resistance (Daborn et al., 2002; Schmidt et al., 2017). Similarly, alleles of the gene acetylcholinesterase (*Ace*) can confer resistance to organophosphate pesticides (Aldridge 1950). Such alleles have been shown to quickly increase in frequency in populations exposed to pesticides (Daborn et al. 2002; Menozzi et al. 2004; Karasov et al. 2010) and in the case of *Ace* and *Cyp6g1*, the resistant alleles are thought to have arisen *de novo* on multiple distinct haplotypes in cosmopolitan populations during adaptation and are therefore often cited as examples of soft sweeps (Karasov et al. 2010; Garud et al. 2015). These results have also been argued to show that adaptation in *D. melanogaster* is not limited by *de novo* mutations (Karasov et al. 2010), but little is known about how the balance of de novo mutations and gene flow has shaped current day patterns of pesticide resistance.

Adaptive introgression results in characteristic genomic signatures that are distinct from both those of neutral introgression and those of classical models of natural selection at the molecular level. First, adaptively introgressed alleles will typically exceed the baseline introgression fraction (Figure 1A). Second, because adaptive haplotypes increase quickly in frequency, the surrounding segments of non-recombined ancestry is expected to be longer than under a neutral model (Shchur et al. 2020). To a first approximation, these patterns are qualitatively similar to classical models of selective sweeps. However, because introgressing haplotypes are genetically distinct and selected alleles are introduced at moderate starting frequencies, the characteristics of genetic variation associated with alleles contributed by adaptive introgression differs substantially (Fraïsse et al. 2014; Racimo et al. 2015; Shchur et al. 2020). Moreover, even neutral admixture affects haplotype patterns, confounding direct quantification of selective coefficients using conventional techniques for selection in single populations (Lohmueller et al. 2011; Racimo et al. 2015). Accurate detection and quantification of adaptive introgression therefore cannot rely on many of the rich and detailed models of adaptive evolution (reviewed for instance in Pavlidis and Alachiotis, 2017) and remains a fundamental challenge in evolutionary genomics (Racimo et al. 2015).

**Figure 1:**
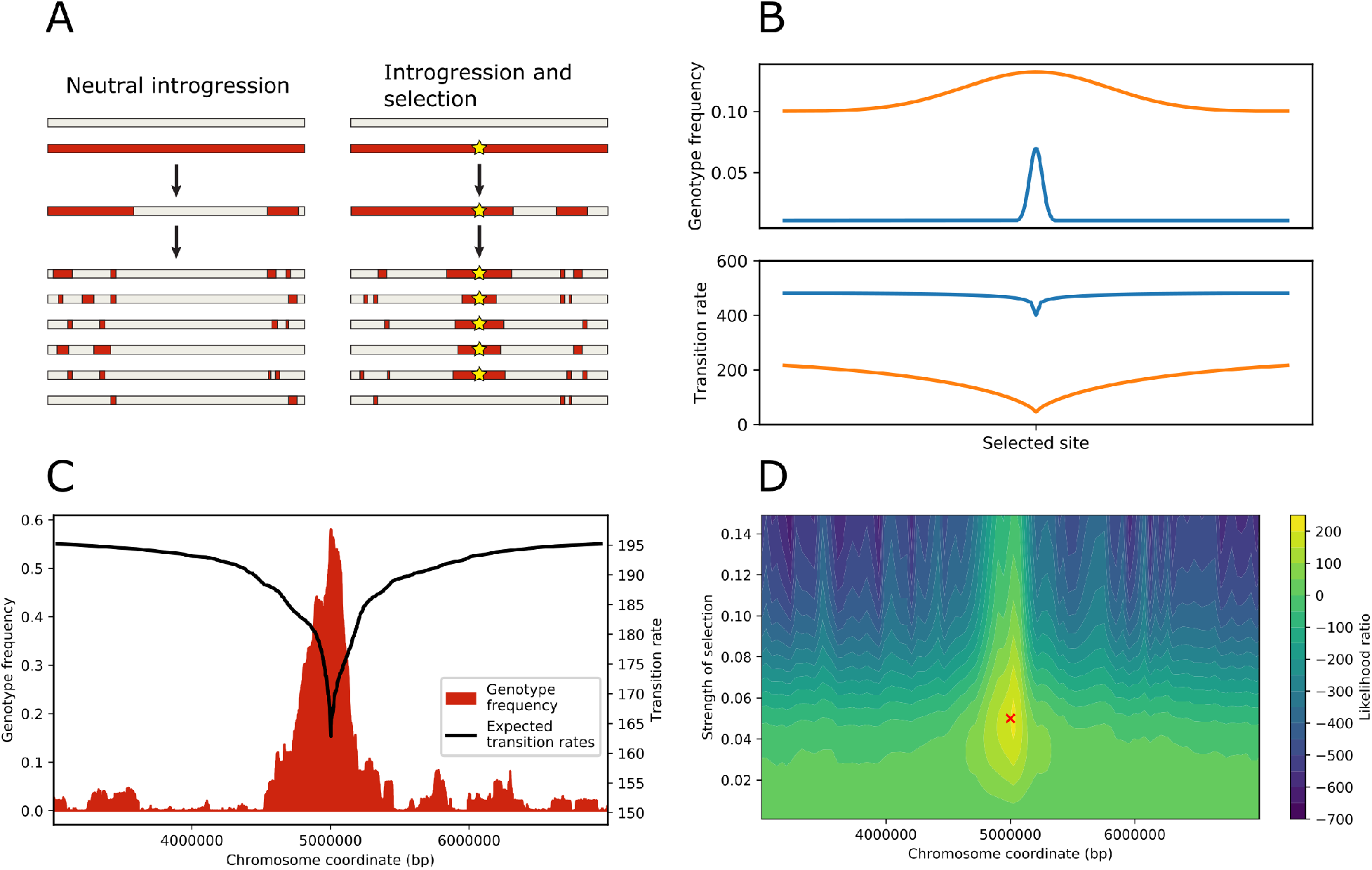
**(A)** Following an admixture event, recombination will break up introgressed haplotypes. In the absence of selection, the frequency of the introgressed genotype (red regions) is expected to remain at a constant low level and haplotype lengths are expected to be short. If positive selection is acting on an introgressed locus (yellow star), the genotype frequency is expected to be higher, and the haplotype lengths larger. **(B)** Cartoon of two scenarios of adaptive introgression. The top panel shows the genotype frequencies of a (blue) 1% introgression pulse of an adaptive locus with weaker selection at position 0, sampled after 400 generations, and (orange) a 10% introgression pulse with stronger selection sampled after 200 generations. The bottom panel shows the corresponding transition rates. **(C)** An example of simulated population with a 1% introgression pulse and a selected allele with s=0.05 at position 5,000,000. The frequency of the introgressed genotype is shown in red, and the expected transition rate of the selected site in black (in transitions from the introgressed genotype to the receiving genotype per Morgan). **(D)** Likelihood surface of simulated chromosome. Adaptive introgression was inferred using AHMM-S for values of s from 0.001 to 0.15 at every 10 sites along the chromosome, for the same simulation as shown in C. The likelihood ratio for each unique combination of site and s is plotted. The red cross marks the position and selective coefficient used in the simulation.

An identifying characteristic of adaptively introgressed alleles is that they reach higher frequencies than neutral alleles (Hedrick 2013; Racimo et al. 2015; Suarez-Gonzalez, Adriana et al. 2018). A first step in specifically searching for adaptive introgression is therefore to infer the ancestry frequencies of admixed samples locally across the genome, which is typically accomplished using Hidden Markov Models (HMM) (Falush et al. 2003; Sankararaman et al. 2008; Baran et al. 2012; Maples et al. 2013). By identifying loci with unusually high proportions of introgressing ancestry within admixed populations, it is sometimes possible to detect signatures of adaptive introgression (Racimo et al. 2015). However, these approaches typically require tailor-made methods for identifying local ancestry outliers consistent with selection. Moreover, as described above, natural selection itself shapes the resulting ancestry tract length distribution, and a more general and powerful method for detecting and quantifying adaptive introgression could explicitly model the consequences of adaptive introgression during local ancestry inference. Recently, a few software packages for detecting adaptive introgression have been released. Genomattn (Gower et al. 2020) uses convolutional neural networks to perform this task, and VolcanoFinder (Setter et al. 2020), can infer adaptively introgressed loci from patterns of elevated heterozygosity surrounding the introgressed allele. VolcanoFinder is intended for mutations that were introduced from a highly divergent population and then went to fixation some time in the past. While it has the advantage of not using data from the donor population, making it applicable in human genetics to detect introgression from unsampled and now extinct hominins, it is less suitable for detecting more recently introgressed adaptive alleles that have still not gone to fixation, or alleles introgressed from a closely related population. The objective of this paper is to develop a method applicable to segregating alleles, from possibly highly related populations, when reference data from the donor population is available.

We have previously developed a method for local ancestry inference (LAI) named Ancestry_HMM (Corbett-Detig and Nielsen, 2017; Medina et al., 2018). Briefly, our approach uses an HMM framework to perform LAI using genomic samples from an admixed focal population and two unadmixed, ancestral reference populations. By assuming a neutral admixture model (Liang and Nielsen 2014), our approach can infer both the timing of multiple admixture pulses and local ancestry patterns across the genome based on low coverage data from unphased diploid samples or samples of arbitrary ploidy, including data generated through pooled sequencing strategies. Because of the generality of this framework, it is both possible and appealing to expand this approach to explicitly model and search for the contributions of adaptive introgression to patterns of local ancestry within samples from admixed populations.

Here, we introduce a novel method called Ancestry_HMM-S (AHMM-S, where S stands for selection) to explicitly model the impacts of natural selection during admixture. Our approach enables the detection of adaptive introgression and estimation of the strength of selection acting on individual loci. We validate this approach through extensive forward simulations, demonstrating that AHMM-S is robust under many plausible scenarios of selection during admixture. We use AHMM-S to analyze a genomic dataset of admixed samples of *D. melanogaster* from a population in South Africa, where we identify several loci that show signatures of adaptive introgression. Our results show that selection has driven cosmopolitan haplotypes carrying insecticide resistant alleles to high frequencies, likely a result of the application of chemical insecticides in South Africa.

## Results and Discussion

### Expected Patterns of Ancestry Transitions During Adaptive Introgression

We began by modifying Ancestry_HMM to estimate the likelihood of adaptive introgression from haplotype patterns surrounding a candidate locus. We did this by adapting a framework for calculating the expected lengths of haplotypes carrying an adaptively introgressed allele that we have previously developed (Shchur et al. 2020), and implemented a fast method for calculating the corresponding transition rates that are used in the HMM. Briefly, we assume a single discrete admixture event, a “one-pulse” model, that took place *t* generations prior to the time of sampling. The probability of the ancestry states in a HMM (emissions probabilities) is not affected by selection and is unchanged from Ancestry_HMM, we therefore refer readers to previous works for details on how these probabilities are calculated (Corbett-Detig and Nielsen, 2017; Medina et al., 2018). However, in order to take selection into account, we must update the transition probabilities to reflect the expected increased frequency of the selected site relative to background levels *i.e*., the initial admixture fraction) (Figure 1B). We do this by modelling the increase in frequency of an additively adaptive allele using a familiar logistic deterministic approximation (Kaplan et al. 1989), as well as the decay of the introgressed haplotypes surrounding the locus through recombination (Shchur et al. 2020) (Figure 1C). By optimizing this model at regular intervals along a chromosome and comparing these results to neutral models, we can detect loci that experience adaptive introgression and quantify the strength of selection that has acted on these sites (Figure 1D).

### Model Evaluation and Validation

In order to validate AHMM-S, we performed forward simulations of selection during admixture. In brief, we simulated admixed populations of diploid individuals which received an introgressive pulse from a second population carrying an adaptive allele, *t* generations prior to sampling. We simulated low coverage, short-read allele counts for 25 diploid individuals, in order to represent a realistic and modest sampling strategy. The simulated reads were conditional on a fixed error rate and their known genotype at each site. We varied the admixture fraction *m* between 0.01 and 0.5 and the selective coefficient *s* of the adaptive allele from 0 to 0.1, and we sampled the population at steps from 50 to 1000 generations. We estimated a selective strength at each locus in our simulated dataset and identified the selected site as the site with the highest likelihood ratio across the chromosome, as might typically be done when searching for adaptive introgression in real datasets.

Our simulations cover an extremely broad range of parameter spaces, and while AHMM-S performs generally well, we were also able to define several important limitations (Figure 2). At lower levels of selection and shorter time periods since introgression, it is difficult to identify the position of the adaptively introgressed locus. The mean distance to the locus is on the order of 1 Mbp, suggesting that the inferred locus is generally incorrect and that outlier values in likelihood ratio is mostly caused by noise. This is consistent with likelihood ratios under these conditions being low (< 50), as well as the inferred selective coefficients being too high. This phenomenon is likely caused by there being little difference in genotype frequency of the selected locus compared to the other neutrally segregating introgressed loci. Since our method depends on there being a difference in genotype frequencies, and correspondingly transition rates, it has low power to distinguish adaptive introgression from noise under these conditions.

**Figure 2:**
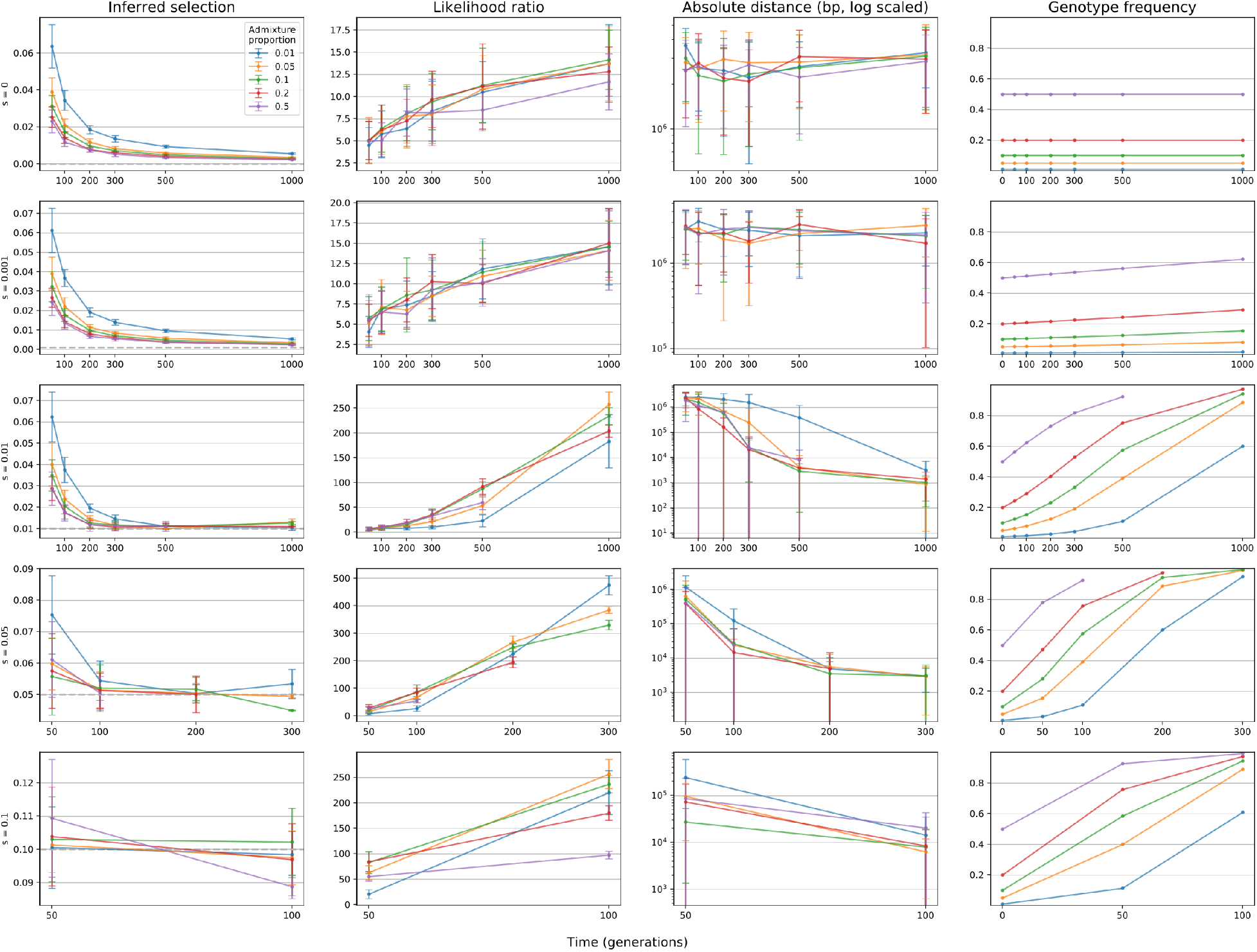
AHMM-S was validated over a range of scenarios for adaptive introgression. The precision of the estimated selective coefficient and genomic location of the selected locus is plotted, together with the corresponding likelihood ratio and frequency of the introgressed genotype. Each row shows a different selective coefficient used in the simulations. Both the likelihood ratio and the precision of the estimates correlate strongly with the frequency of the selected locus. The x-axis shows the sampling time in generations since admixture. Each datapoint shows the mean value of 20 simulations, and error bars show the standard deviation.

Nonetheless, the method performs well in general for a range in the plausible parameter space, and we find that for populations sampled 300 or more generations after admixture with moderate selection (s=0.01), the model accurately estimates both the position of the selected site and the strength of selection. In cases of stronger selection (s > 0.05), the inferred position and estimated selective coefficient are close to the real value already at 50 or 100 generations. Under favorable conditions, we are able to identify the correct position within a few kbp, which is close to the limit of resolution in our simulations, which are based on SNP densities found in *D. melanogaster*. The inferred selective coefficients are also close to the real value under these conditions, and the error is within 20%.

Overall, AHMM-S works best when selection is strong and/or when selected sites have reached frequencies that are significantly higher than baseline introgression levels. These are conditions that are most likely to be associated with important phenotypic changes and are for this reason of greatest interest to biologists. Nevertheless, weakly selected sites may reach high frequencies over longer periods of time. Unfortunately, this also means that recombination will have time to break diagnostic haplotype patterns apart, decreasing the ability to quantify the relevant selective coefficient. This pattern is expected over time scales beyond what we validate for here and that is much longer than is suitable for most LAI applications.

### Effects of algorithm for calculating expected transition rates

We evaluated the simulated scenarios above using the four-point analytical approximation of the expected transition rates. This method calculates expected transition rates for four sites spaced along the chromosome and then interpolates the rates for all other sites, but we also evaluated the performance of a slower forward iteration-based approach, where the expected transition rate is calculated between each pair of loci along a chromosome for a given model of selection. We applied the same simulation strategy for a subset of population parameters using the forward iteration algorithm and inferred adaptive introgression (Figure S1). Both methods perform similarly well across a range of input parameters, and we therefore decided to only use the four-point approximative method for further analysis.

### Effects of sample size and sequencing approach

We tested the effects of sample size by increasing the number of individuals in the set of simulated reads from 25 to 75 (Figure 3). While a larger sample size improved the estimates of both the selective coefficient and the location of the locus, the effects on the estimated selection coefficient was generally small. Therefore researchers should use the largest feasible sample size when studying adaptive introgression using this method, although the primary results from the modest sample sizes suggest that this method is applicable for all species but those that are most challenging to sample.

**Figure 3:**
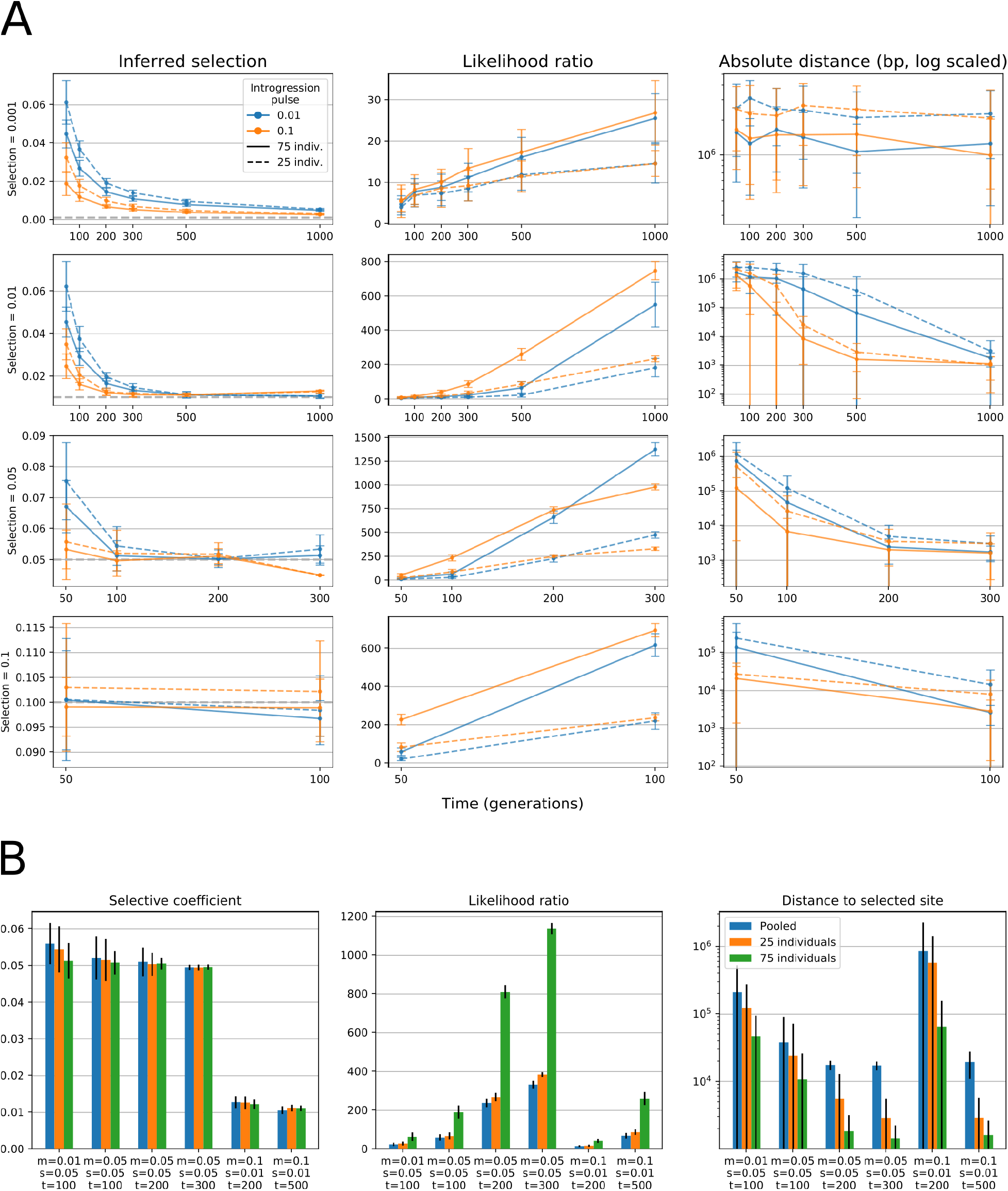
(A) Comparison of precision of estimated selective coefficient and positions for 25 and 75 individual samples. Increasing the sample size improves the estimate of both s and p, but has a stronger effect on the precision of the inferred position. (B) Effects of sampling strategy on inference of adaptive introgression. Forward simulations of adaptive introgression were converted to simulated reads of either 25 or 75 diploid individuals sampled separately, or of 25 diploid individuals sampled as a pooled set of reads. The effects on the inferred selective coefficient are minor, but increased sampling improves the inference of the location of the selected site.

We also investigated the performance of AHMM-S when using pooled sequencing data. A subset of the simulated populations were converted to pooled reads (instead of reads separated by individual), and adaptive introgression was then inferred (Figure 3B). The estimated strength of selection is close to the real value, and errors ranging from 5 to 25%, which is comparable to the values generated when using individually sequenced samples instead. The accuracy of the position estimate is worse than for individually sequenced samples, though it also improves with stronger selection and longer time since admixture. We therefore suggest using individually barcoded and sequenced samples, but pooling may provide an economic tradeoff if accurately mapping the specific selected site is not a primary concern.

### Generating a null model

AHMM-S performs a likelihood test for each site (or a subset of sites) on a chromosome, and for relatively densely spaced markers this implies a substantial number of hypotheses are tested in a single run of the program. As many of the sites will be in genetic linkage with each other, these tests are not independent. Furthermore, since linkage is expected to decay over time, identifying a cutoff for statistical significance is difficult and to some degree depends on the time since admixture. For this reason we recommend performing simulations of a neutral introgression scenario with similar population parameters as the dataset of interest. The distribution of likelihood ratio scores generated in these simulations can then provide a null model for variation in likelihood scores under a neutral admixture model.

### Computational performance

The computational performance of AHMM-S is influenced by the number of sites, the number of samples and the type of algorithm used for approximating the trajectory of a selected allele. A dataset with 20,000 sites and 25 diploid individuals takes approximately 2 hours using a single thread of an Intel i7-8550U CPU when using the 4-point approximation method for computing the transition rates and 6.5 hours for the same dataset when using the forward iteration method to obtain transition rates. Memory usage is similar for both methods with 120-140 Mb used for both algorithms. When using a larger dataset consisting of 75 individuals, computation time is approximately 4 times longer, and memory usage rises to 700 Mb.

### Robustness to Parameter Misspecification

AHMM-S assumes knowledge of several demographic parameters, including the time of introgression, the admixture fraction and the effective size of the admixed population. In practice, these must also be estimated from the data and the true parameters cannot be known without some uncertainty. We have previously shown that local ancestry inference using Ancestry_HMM is neither strongly affected by parameter misspecification nor the presence of selection (Corbett-Detig and Nielsen 2017), but that the estimated times since introgression were. We therefore evaluated the robustness of estimates of selective strength by AHMM-S and the consequences of poorly estimated parameter values, by intentionally misspecifying the necessary parameters on a subset of the simulations used for validation. While misspecifying population size has little effect on the final estimated selective coefficient, both time since introgression and the admixture fraction can skew the estimate (Figure S2). Even so, if the specified values are within 20% of the true values, the errors in estimation of *s* are within 40%.

In general, it is straightforward to accurately estimate the overall admixture fraction using a range of approaches (for instance Alexander et al., 2009; Pritchard et al., 2000) and our previous work has shown the time of admixture can be approximated well using approaches that we (Corbett-Detig and Nielsen 2017; Medina et al. 2018) and others (Pool and Nielsen 2009; Gravel 2012; Loh et al. 2013) have developed, even with moderate impacts of natural selection. In contrast, the effective sizes of admixed populations may be challenging to accurately infer. However misspecification of the effective population size has only a minor impact on estimates of selection obtained using this method, suggesting that our approach is robust even to substantial uncertainty regarding the effective population size.

### Effects of continuous gene flow, going to fixation, dominant/recessive selection and segregation in the donor population

AHMM-S uses a simple model for adaptive introgression, which assumes 1) a single pulse of introgression, 2) that the selected allele is fixed in the donor population, and 3) that selection is additive for the adaptively introgressed allele. We tested the performance when violating these assumptions by simulating populations where there was either continuous gene flow for 20-100 generations, where the selected locus was segregating at 50% in one ancestral population or where the selected allele was either dominant or recessive. The results are summarized in Figure S3–S5. In general, AHMM-S is able to identify adaptive introgression in these cases, but with somewhat reduced precision compared to when the population model is not violated.

A further limitation of AHMM-S is that it is not capable of estimating the selective coefficient when the selected allele has gone to fixation. This is caused by the method that is used to calculate the expected transition rates (Figure S6). In such a scenario, the program is still generally capable of identifying the location of the adaptively introgressed site, but the reported value of *s* will be lower than the true value. As it is possible to quantify the local ancestry along the chromosome with the software package on which AHMM-S is based (Corbett-Detig and Nielsen 2017), it should be easy to identify such cases.

### The effects of a small introgression fraction

AHMM-S calculates expected transition rates based on a logistic function for calculating the allele frequency trajectory of the selected site. As noted in (Shchur et al. 2020) this logistic function is only a good approximation of the trajectory when the admixture pulse is large. Shchur et al. developed a simple stochastic method based on multiple forward simulations for estimating the trajectory when the admixture pulse is small, and we implemented it in AHMM-S. We compared the logistic and the stochastic methods for simulations of populations with m=0.001 and m=0.0001, where we conditioned the simulations on the selected allele not being lost by drift. In our simulations (Figure S7), we see little effect of a small m on our estimates, and no large difference between the logistic and stochastic methods for approximating the trajectory of the adaptive allele. As the logistic approximation is significantly faster, we would recommend its use even in cases where the initial admixture pulse is very small.

### Linkage between multiple selected sites

Multiple selected sites can be located near each other, and we examined how AHMM-S can handle such a scenario and how it affects inference of selective coefficients. We ran simulations where two positively selected sites were placed at varying distances from each other, ranging from 0.1 to 5 cM, and inferred adaptive introgression (Figure S8). When the two sites are located in close proximity (0.1 cM), AHMM-S will typically produce a single peak for both the likelihood ratio and the inferred selection. The inferred selection coefficient is close to the sum of *s* for each site, as would be expected for additive selection. When the sites are placed at increasingly large distances, individual peaks are distinguishable from 1-2 cM, and the additive effect is diminished as the distance increases. Care must still be taken though, when interpreting the inferred selective coefficients in such cases as they are likely to be overestimations reflecting in part the joint effect of two sites.

It is also possible that negatively selected loci affect the inference of adaptive introgression at linked sites. Purging of weakly deleterious alleles following admixture has been suggested in a number of systems (Harris and Nielsen 2016; Kim et al. 2018; Meiklejohn et al. 2018) often causing large changes in introgression patterns along the genome. Although we do not consider this effect here, we expect that patterns of large peaks of introgressing ancestry are unlikely to occur under a model of purely weakly deleterious variation and is unlikely to dramatically affect inferences using this method.

### Detecting negative selection

AHMM-S can detect negatively selected sites as well by treating negative selection of the introgressed genotype as positive selection of the receiving genotype (defined as the less common genotype and more common genotype respectively). To test how well this works, we ran a smaller set of simulations (s was set to −0.05 and m to either 0.1 or 0.01) where a single introgressed allele experienced negative selection following admixture. Figure S9 shows the case where m is 0.1, and while we can easily detect the position of the negatively selected locus, the introgressed allele will quickly be lost, which leads to an underestimate of the selective coefficient. For this reason, we expect that it will often be challenging to detect negatively selected alleles with small introgression fractions. However, if selection is weak and the introgression pulse is large the allele will segregate for a significant time and AHMM-S will be able to generate a reasonably accurate estimate of the selective coefficient.

### Suitability for detecting Dobzhansky-Muller incompatibilities

A specific example of negative selection that is expected to be common following admixture between species and distantly related populations are Dobzhansky-Muller Incompatibilities (DMIs) (Coyne and Orr 2004) and there has also been considerable interest in identifying DMIs within admixed populations (Corbett-Detig et al. 2013; Schumer et al. 2014; Pool 2015; Powell et al. 2020). DMIs are caused by complex epistatic interactions between at least two different loci and we therefore evaluated how our approach might cope with such scenarios. As a proof of concept, we simulated several two-locus DMI scenarios and used AHMM-S to identify sites which show signs of selection. We found that our method consistently identifies selected sites with high accuracy, i.e. selected loci are detected within 8 kbp. As expected, due to the conditional nature of selection against DMI loci, estimated selective coefficients are typically small relative to a single locus model, with estimates 50-90% lower than the actual value (Figure S10). Our approach may therefore be applicable for detecting DMI’s in addition to adaptive alleles. However, we caution that without additional evidence (*e.g*., linkage disequilibrium in admixed samples) or experimentation to demonstrate epistatic selection, it will not typically be possible to distinguish between DMI’s and strong single locus selection based purely on the results from our program. If possible, putatively selected sites should be biologically interrogated to identify specific likely modes of selection. Furthermore, parameters such as the distance between the incompatible loci, the size and timing on the admixture pulse can affect the strength and direction of selection on incompatible loci, and comparing real data to simulations with similar parameters may also provide increased confidence in the presence of DMIs.

### *Adaptive Introgression in* D. melanogaster

In order to test AHMM-S on real data we selected a population sample of *D. melanogaster* from South Africa that has shown signals of admixture in previous studies (Lack et al. 2015; Corbett-Detig and Nielsen 2017; Medina et al. 2018). This dataset is moderately sized (n=81), the admixture history is approximately consistent with a one-pulse admixture model, and the previously estimated time since admixture (m=0.17, t=430 generations) suggests that this population is ideally suited for testing our approach (Corbett-Detig and Nielsen 2017; Medina et al. 2018). First we performed simulations of neutral admixture in a similar population, to determine the null model against which we test for adaptive introgression. We identified likelihood ratio outlier peaks (Methods) and then determined the likelihood ratio threshold that would generate on average a single false discovery per genome. In our case, this threshold is 15; i.e. we expect one likelihood ratio outlier above 15 under a neutral model.

We then applied our method to this population, where we observed highly variable patterns of adaptive introgression across the genome. Specifically, we identified one locus on chromosome 2L, three on 2R, 13 on 3R and none on 3L and X as putative targets of selection following admixture, with selection coefficients ranging from 0.0046 to 0.0115 (Figure 4, Table 1). We therefore find evidence for moderately strong fitness effects associated with introgressing cosmopolitan ancestry in the focal population. The inferred selective coefficients are consistent with our expectations given the relatively short time since admixture, in which selection would need to be relatively strong to drive alleles to moderate frequencies.

**Figure 4:**
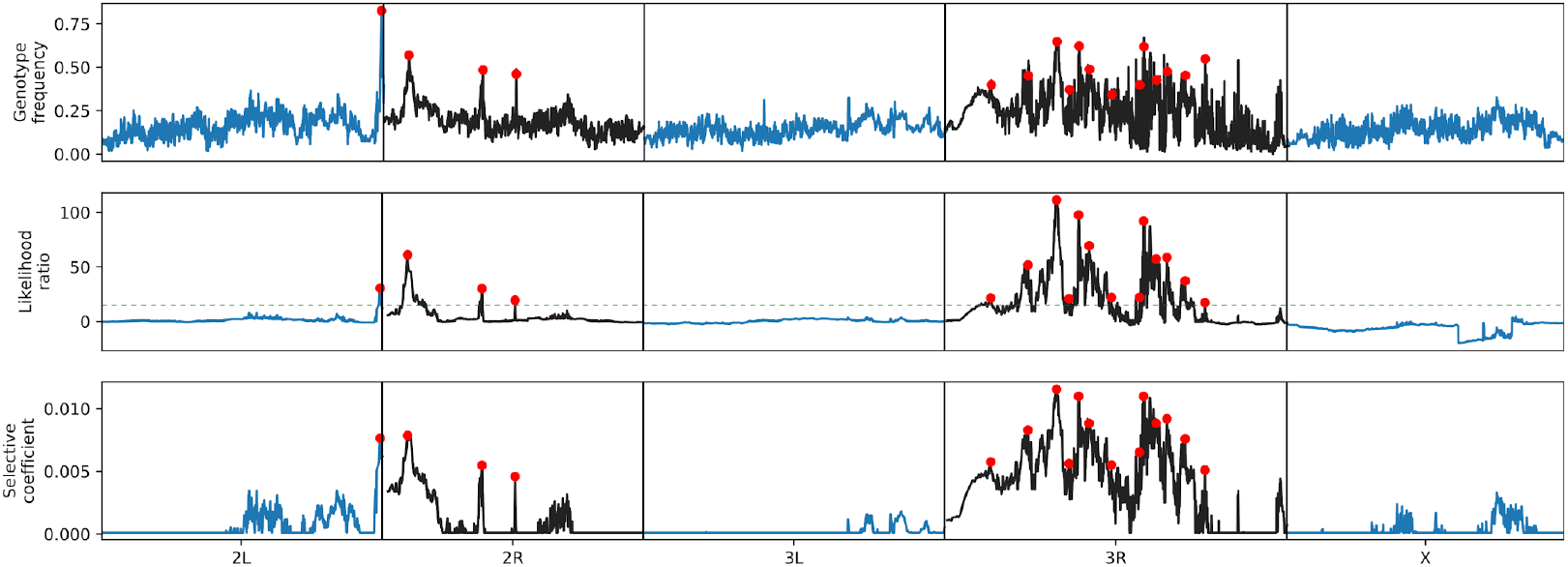
*Signals of introgression across the genome in a South African population of* D. melanogaster. *17 loci (red dots) showed evidence for adaptive introgression (likelihood ratio > 15, dotted green line) and were located with a distance between peaks of at least 2 cM. Most putatively selected loci are located on chromosome 3R. There’s a clear correlation between the frequency of the introgressed genotype (top panel) and the likelihood ratio (middle panel) and inferred selective coefficient (bottom panel*).

**Table 1:** Candidate loci for adaptive introgression in South African *D. melanogaster*

| Chromosome | Position (bp) | Frequency of introgressed genotype | Estimated Selective coefficient | Likelihood ratio | Candidate genes (within 5kbp) |
| --- | --- | --- | --- | --- | --- |
| 2L | 22782051 | 0.83 | 0.0076 | 30.5 |  |
| 2R | 6174331 | 0.57 | 0.0079 | 61.1 | tRNA:Arg-ACG-1-3, tRNA:Lys-CTT-1-5, tRNA:Arg-ACG-1-4, Cyp6w1, tRNA:Arg-ACG-1-5 |
| 2R | 12182790 | 0.48 | 0.0055 | 30.0 | EndoG, asRNA:CR45264, CG8860, wash, CG33964, CG13175, SmF, Cyp6g1 |
| 2R | 14869645 | 0.46 | 0.0046 | 19.4 | Pcf11, Cyp6a22, Cyp6a17 |
| 3R | 13250975 | 0.65 | 0.0115 | 111.4 | Ace |
| 3R | 15037772 | 0.62 | 0.0110 | 97.8 | RpL10Aa, lncRNA:CR44944 |
| 3R | 20295870 | 0.62 | 0.0110 | 92.2 | cic |
| 3R | 15887044 | 0.49 | 0.0088 | 69.4 | CG14877 |
| 3R | 22181404 | 0.48 | 0.0092 | 58.7 | SKIP |
| 3R | 21318439 | 0.43 | 0.0088 | 57.2 | lncRNA:Hsromega, mir-4951, CG16791 |
| 3R | 10929354 | 0.45 | 0.0083 | 51.8 |  |
| 3R | 23666552 | 0.45 | 0.0076 | 37.2 | CG31145 |
| 3R | 19962983 | 0.40 | 0.0065 | 22.0 | Hs6st |
| 3R | 17696124 | 0.34 | 0.0055 | 21.9 | lncRNA:CR46036, osa |
| 3R | 7919356 | 0.40 | 0.0057 | 21.5 | CG2678, CG10445, lds,<br>scaRNA:MeU2-C41, dsx |
| 3R | 14266513 | 0.37 | 0.0056 | 20.9 | CG14841, CG14839, trx,<br>asRNA:CR46020 |
| 3R | 25262165 | 0.55 | 0.0051 | 17.3 | Nup358, CG11857, CG10425,<br>asRNA:CR46099, CG11858,<br>GlnRS, RIOK2, asRNA:CR45214 |

Selection for resistance to commonly used insecticides might underlie many of the signatures of adaptive introgression that we observe. Several candidate loci are located close to genes known to be associated with resistance, such as the three loci on 2R, which are all located within 5 kbp from *Cyp6* Cytochrome 450 genes. Two of the most prominent likelihood ratio outliers are located next to *Cyp6g1* and *Cyp6w1* respectively, and allelic variants of these genes are specifically known to confer resistance to DDT exposure (Daborn et al. 2002; Schmidt et al. 2017). The third candidate locus is located close to a cluster of several *Cyp6* genes, of which *Cyp6a17* and *Cyp6a23* have been shown to be associated with resistance to other insecticides (Battlay et al. 2018). Additionally, the candidate site with the highest likelihood ratio on chromosome 3R is located within the acetylcholinesterase (*Ace*) gene. Several different common alleles of *Ace* confer resistance to the large class of organophosphate insecticides and it is a locus that is known to be under selection when these insecticides are introduced (Karasov et al. 2010; Garud et al. 2015). Resistance to insecticides is a strong candidate phenotype driving adaptive introgression because DDT and most other insecticides were first applied for insect control in populations outside of Africa, where resistance originally evolved (Schmidt et al. 2017). DDT is still actively used in South Africa to control mosquito populations (Biscoe et al. 2005), and the country imports a wide range of other broad-spectrum insecticides (Quinn et al. 2011). Our results therefore strongly suggest that resistance to commonly used pesticides has been a primary driver of adaptive introgression in admixed populations of *D. melanogaster*.

In total, 17 loci are classified as potential candidates for adaptive introgression. We performed a gene ontology (GO) analysis of genes that were either spanning, or were located within 5 kbp of the candidate locus (N=46). Two categories, “organic cyclic compound binding” and “heterocyclic compound binding” (both N=21) showed a significant enrichment (at q<0.1) after correcting for false discovery rate (Table S1). We note that these GO categories contain many more genes in addition to *Ace* and *Cyp6* genes, presenting the possibility that our method may have identified genes that contribute to insecticide resistance in nature and that were not previously known (Table 1, S1). For instance, of the other potential adaptively introgressed genes, *lncRNA:Hsrω* has been shown to provide some protection against insecticides in laboratory experiments (Chowdhuri et al. 2001), but it has not been identified in selection scans. As each candidate locus is located close to several genes, further functional work is necessary to determine exactly which genes are driving the signatures of selection and their specific phenotypic effects.

Our findings here mirror earlier studies on selection in *D. melanogaster*, and two of our most distinct outliers, *Ace* and *Cyp6g1*, have also been strong outliers in selection scans presented in previous papers (*e.g*., Garud et al., 2015; Karasov et al., 2010). In the case of *Ace*, several different alleles which confer resistance to insecticides have been found at high frequencies in cosmopolitan populations, and Karasov et al. argues that this suggests that adaptation in this species is not limited by the mutation rate. These mutations are thought to have appeared in cosmopolitan populations and loci showing signs of adaptive introgression in our dataset is consistent with this idea, especially since the majority of *D. melanogaster* genetic diversity is found in Africa (Begun and Aquadro 1992; Thornton and Andolfatto 2006; Pool et al. 2012). On the other hand, this suggests that adaptation to pesticides is not driven by additional *de novo* mutations in our dataset, but instead it is shaped in large part by introgression.

In Garud et al., three genes showed the strongest signals for recent strong selection. Besides *Ace* and *Cyp6g1*, which we find here, *CHKov1* was also a major outlier. A transposon insertion inside *CHKov1* is associated with resistance to both infection by the sigma virus and to organophosphate pesticides (Aminetzach et al. 2005; Magwire et al. 2011). We do not find *CHKov1* among our set of genes, but it is located approximately 60 kbp from one candidate peak (on chromosome 3R position 25262165). It is possible that we were not able to identify the exact position of this gene with sufficient precision, but simulations above suggest that we can accurately map loci with this strength of selection. It is also possible that cosmopolitan alleles at this locus are not selected within our focal population. Consistent with this idea, *CHKov1* is thought to have spread from standing variation that was present within the ancestral African populations (Aminetzach et al. 2005; Magwire et al. 2011). Our results are therefore concordant with expectations from known geographic distributions of strongly selected insecticide resistance loci and further support the idea that resistance to insecticides has been an important driver of adaptive introgression.

### Conclusions

Generalized tools for inferring and mapping adaptive introgression and estimating the strength of selection from genomic data have long been lacking. Here we provide an approach for this problem that is well-suited for detecting adaptive introgression in commonly generated datasets. AHMM-S can both identify the locations of adaptive introgression genes and infer their selective coefficients. It is robust over a range of introgressive scenarios, and especially in cases where an adaptively introgressed gene has strongly increased in frequency. Such scenarios are especially important, as the loci with the most severe shift in genotype frequencies are more likely to be important for understanding adaptation. Many previous studies have addressed the question of adaptive introgression by tailoring analysis methods to the specific data at hand (for instance Sankararaman et al., 2014; Vernot and Akey, 2014), but AHMM-S provides a more general solution and enables searches for adaptive gene flow over a large range of eukaryotic species.

AHMM-S is based on a simple model of adaptive introgression with positive selection of a single locus following a single introgression pulse, but can still identify adaptive introgression with reasonable precision from more complex scenarios, such as continuous gene flow. To improve the performance by explicitly model these phenomena when generating the expected transition rates. We could model negative background selection around a positively selected site, or different selective regimes such as balancing or conditional selection, to improve our estimates of selective strengths. In the case of balancing selection, we would expect to identify the selected locus (given the introgression pulse is different from the equilibrium frequency of the selected site), but it would be interpreted as a positively selected site. For these more complex scenarios, such as balancing selection or epistatic interactions, it will most likely be easiest to compare selective landscape to the landscape generated from simulations to infer the exact nature of the adaptive evolution that has taken place.

In this work, we also use AHMM-S to investigate possible adaptive introgression in *D. melanogaster* in South Africa. Several of the loci we identify are associated both with pesticide resistance and with strong selection in cosmopolitan populations, suggesting that pesticide use has been a major driver of selection both in sub-Saharan Africa and elsewhere. Though where some of these resistant alleles are thought to have evolved through *de novo* mutations in cosmopolitan populations, we here see that they have been introduced through introgression in South Africa. This positions admixture to be an important factor to consider when studying the causes of adaptation and selection within species and populations.

## Materials and Methods

We developed an approach based on an adaptation of Ancestry_HMM (Corbett-Detig and Nielsen 2017; Medina et al. 2018) that allows one to infer adaptive introgression by implementing a model for tract length distributions surrounding an adaptively introgressed locus (Shchur et al. 2020). This allows us to calculate expected transition rates between the two genotypes at a given distance from a locus of interest, which then can be used in a HMM that can calculate a likelihood score for a particular value of *s* at a particular site. We assume a single discrete admixture event, a “one-pulse” model, that took place *t* generations prior to the time of sampling. Therefore, the state space of the HMM is all possible counts of chromosomes of ancestry type one given the number of chromosomes, or ploidy, of a sample (*e.g*., for a diploid, H = {0,1,2}). The probability of the ancestry states at a given site in an admixed genome is unchanged in our modified framework (emissions probabilities, see (Corbett-Detig and Nielsen 2017; Medina et al. 2018)). However, to incorporate natural selection, we must update the transition probabilities to reflect the increased frequency of the selected site relative to background level. We define a three locus coalescent process where one site denotes an introgressing allele experiencing additive selection. The other two sites trace the ancestry linked to that site. For example, at the time of admixture, the only possible three locus haplotypes are 0*-0-0 and 1*-1-1, where * denotes the selected locus. By tracking the frequencies of recombinant haplotypes, i.e. chromosomes in which the two linked sites correspond to different ancestry states (0*-1-0, 0*-0-1, 1*-1-0, 1*-0-1), we can define an ancestry transition model along the chromosome in the regions adjacent to the selected site. Given a dataset with known recombination rates between each site, AHMM-S can then generate expected transition rates going away from that site in each direction, either by using a forward iteration strategy, where the transition rates between adjacent sites are calculated for each site along a chromosome, or through an approximative method which can interpolate transition rates based on just four sites.

When running AHMM-S, a single chromosome dataset is specified, together with the population size *N*, introgression fraction *m* and time since introgression *t*, which all need to have been estimated previously. AHMM-S will then estimate a likelihood ratio for a specific site *p* and a specific selective coefficient *s* compared to the neutral case for that site. AHMM-S will loop over all sites, or a subset of sites, and can then either calculate the likelihood ratio in a grid for a defined set of values of *s*, or find the value of *s* that gives the highest likelihood for each site, by using golden section search. The size of window used in the HMM can be specified by the user, as can the number of sites that will be analyzed.

### 4-point approximation

The transition rates between ancestries of different types are functions of recombination distance *r* from the site under selection. In particular, the transition rate *f*_10_(*r*) from ancestry type 1 to ancestry type 0 is a monotonously growing function with a finite limit at infinity which is equal to the transition rate under neutrality. Similarly, the transition rate *f*_01_ (*r*) is a monotonously decreasing function. We will search for an approximate solution of these functions in a form of

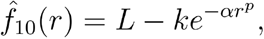

where coefficients *L*, *k*, *α* and *p* can be calculated as follows. Let *r*_1_ and *r*_2_ be two points such that 0 < *r*_1_ < *r*_2_ < ∞. We can estimate transition rates numerically for *r* = 0, *r*_1_ and *r*_2_.

In order to be informative, *r_2_* can be set to the value of expected tract length under neutrality, and we set *r*_1_ = *r*_2_/10. The expected tract length under neutrality can be calculated analytically under SMC’ model (Marjoram and Wall 2006). Using the formula derived by (Liang and Nielsen 2014), we set

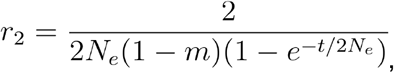

where *N_e_* is the effective size of the admixed population, *m* is the admixture fraction and *t* is the time of introgression.

Given these values, we can calculate the coefficients of the function 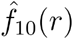. *L* is the transition rate for the neutrality (*s* = 0), because 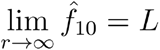. Again, following (Liang and Nielsen, 2014), the neutral transition rate is given by the following formula

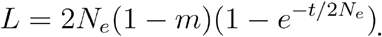

Next, for *r* = 0 we have 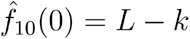, hence

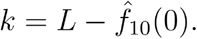

In order to find the last two parameters *α* and *p*, there are two more equations corresponding to *r* = *r*_1_ and *r* = *r*_2_. Denote

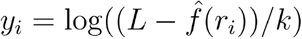

for *i* = 1, 2. Then

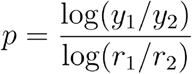

and

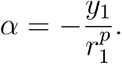

We validated this approximation by comparing with simulations and with “forward time” approximation, see Figure S11 for an example. In all the considered scenarios, our approximation turned out to be very precise.

### Simulations

We validated the method using extensive simulations of populations of 100,000 diploid individuals over a range of parameter values. We varied the selective coefficient *s* from 0 to 0.1 (0, 0.001, 0.01, 0.05, 0.1) and the admixture fraction *m* from 0.01 to 0.5 (0.01, 0.05, 0.1, 0.2, 0.5). We let the simulations run to *t* = 1000 generations or 0.99 frequency of the selected allele, whichever came sooner, and we drew samples from the admixed population at 50, 100, 200, 300, 500 and 1000 generations. 20 simulations were run for each parameter set and 25-75 diploid individuals were sampled per time point per simulation. The selective coefficient s is specified for the diploid case, and selection acts additively, meaning that a heterozygous individual experiences half the selective strength.

Simulations were performed using SELAM (Corbett-Detig and Jones 2016), which is a forward simulator that records the full local ancestry across the genome. We then converted the haplotype information generated by SELAM to simulated genotypes representing the reference panels of the two ancestral populations, using the results of a coalescent simulation consistent with the evolutionary history of ancestral *D. melanogaster* populations (following Corbett-Detig and Nielsen, 2017; Pool et al., 2012). Specifically, we used the SMC’ coalescent simulator, MaCS (Chen et al. 2009) with the following command line:

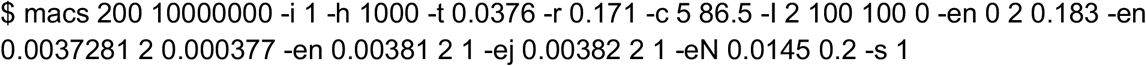

To construct admixed individuals, we applied genotypes obtained from each MaCS-simulated population to fill in genotype data on the appropriate chromosomes for each ancestry type. Pairs of chromosomes were joined to create single diploid individuals, and then we simulated shortread pileup data for each variable site by drawing the depth from a Poisson distribution with mean 2 and the resulting alleles from a binomial distribution with *p* equal to the genotype frequency in the individual sampled at that site and applying a uniform error rate of 0.01 per allele per site.

We emphasize that users interested in evaluating the utility of this or our previous software packages can produce similar simulations tailored to their specific evolutionary models or sequencing and sampling schemes by applying our method using the LAI simulation packages released in (Schumer et al. 2020).

We then analyzed the simulated reads with AHMM-S, and specified the values of *N*, *m* and *t* that were used in each simulation. We analyzed either 25 or 75 diploid individuals, or 50 pooled chromosomes, depending on experiment. We used the 4-point approximative method for calculating transition rates, and included sites in the HMM in a window extending 10% of the chromosome length in each direction of the focal site. We used the golden section search algorithm and extracted the site that had the highest likelihood ratio value as the inferred adaptively introgressed site. For the pooled samples, only every 100 sites were analyzed, in order to reduce computational time.

### Robustness

The robustness of AHMM-S against misspecification of input parameter values was evaluated using a subset of the same simulations that were generated for validation. When running AHMM-S, we varied population size (10,000 and 1,000,000 instead of 100,000), introgression fraction *m* and introgression time *t*, setting them to 10% and 20% above and below the true value. Each parameter set was run in 20 replicates.

### Simulations for testing the effects of model violations

We performed simulations to test the effects of continuous gene flow, segregation in donor population, dominance and fixation in focal population on performance of AHMM-S in estimating position and selective coefficient. All simulations were performed in 20 replicates. Introgression pulse was set to 0.01 or 0.1, selective coefficient to 0.01 or 0.05. The simulations were performed as described above, with the following differences: for continuous gene flow, we set an initial introgression pulse to 0.01 or 0.1, and then allowed for 1% gene flow for 20 generations or 0.2% gene flow for 100 generations. When simulating an adaptive allele segregating in the donor population, we set the allele frequency to 50% at the time of introgression. To simulate dominant or recessive selection, we modified the SELAM selection file to change selection to work in a completely dominant or recessive way.

### Simulating a small admixture pulse

When running simulations with a small admixture pulse (m=0.001 or 0.0001), there is a strong risk that the adaptively introgressed site would be lost to drift. In order to counteract this, we condition our simulations on not being lost, by restarting SELAM if the site was lost, until we had 20 replicates for each set of parameters. After this the AHMM-S pipeline was ran as usual

### Multiple selected sites

We tested the ability of AHMM-S to distinguish adaptively introgressed sites that are located near each other by running simulations as described above, but with two selected sites with equal selective strengths at 0.1, 1, 2, and 5 cM distance from each other. Three parameter sets were used: introgression fraction *m*=0.01, selection *s*=0.025 and time *t*=500 generations; *m*=0.1, *s*=0.005 and *t*=200; and *m*=0.17, *s*=0.005 and *t*=430. The last parameter set corresponds to the population of *D. melanogaster* analyzed in this study. We then analyzed the simulations with AHMM-S and plotted the inferred selective coefficients and likelihood ratios to determine peak separation.

### Detecting negative selection

Simulations of introgression followed by negative selection of an introgressed allele located in the middle of the chromosome was performed using SELAM as previously described. AHMM-S is designed to infer positive selection on the introgressed genotype, and in order to make it detect negative selection, we converted the genotype identity of the simulated individuals, so negative selection of the introgressed genotype is interpreted as positive selection of the receiving genotype. We then converted the simulation files to reads as before and analyzed it with AHMM-S.

### Dobzhansky-Muller incompatibilities

Simulations of diploid individuals were performed as described above for two different scenarios of DMIs. In both cases, two loci (A and B) had a negative DMI interaction. At each locus two alleles were present (A0 and A1, B0 and B1) where the integer denotes the ancestral population that contributed the relevant allele. In scenario one, an individual carrying any combination of alleles that were not from a single population (i.e. A0+B1, or A1 +B0) had a reduced fitness (s<1). This is an example of dominant sign epistasis. In scenario two, only one specific combination (*i.e*., A0+B1 but not A1+B0) was selected against (s<1). In both cases all other allelic combinations had s=1. We ran these scenarios for three values of *s*: −0.01, −0.05 and - 0.1, with a 50% introgression pulse. The two sites were placed on the same chromosome at a distance of 40 cM, where linkage plays little role in governing outcomes. We then converted the simulations to reads as previously described and analyzed them using AHMM-S. Since DMIs cause negative selection at the interacting loci, and AHMM-S is only designed to handle positive selection, we changed genotype identities before converting the simulations, which allowed us to treat the negative selection at these loci as positive selection and bypass this limitation.

### D. melanogaster *data*

We applied AHMM-S to a publicly available dataset of *D. melanogaster* from South Africa and Europe (Lack et al. 2015, Lack et al 2016). We extracted both homozygous and heterozygous regions for each sample and ran AHMM-S after supplying the appropriate ploidy for each (*i.e*., 1 if inbred, 2 if outbred) as we have done previously (Medina et al. 2018). We also removed any chromosome arm containing one of the common chromosomal inversions in this species (Corbett-Detig and Hartl 2012; Lack et al. 2016). As these assemblies are generally quite high quality, we used the genotype emissions function in AHMM to calculate the local ancestry across the genome (Corbett-Detig and Nielsen 2017). We also supplied a fine-scale recombination map for this species (Comeron et al. 2012).

Because the direction of introgression is known, *i.e*., populations with cosmopolitan ancestry have recently back-migrated into Africa, we scanned specifically for adaptive introgression of cosmopolitan alleles into these predominantly African populations. We performed 50 simulations of a neutral admixture scenario with the same population parameters (m=0.17, t=430, N=100000) and used these to determine a likelihood ratio cutoff that would produce an acceptable false positive rate. A likelihood ratio > 15 and and filtering for proximity to other higher peaks produced 42 outliers above this threshold, corresponding to a false positive rate of ~5% in the D. melanogaster dataset. Using simulations, we also determined that we could distinguish selected sites separated by at least 2 cM (see section on multiple sites). A GO enrichment analysis was performed on the candidate loci using Gowinda (Kofler and Schlötterer 2012). We ran the program with default parameters and separately included only genes that either spanned the candidate locus or included genes located 5 kbp upstream or downstream of the locus.

## Contributions

JS developed the software, performed simulations and validation, and analyzed the *D. melanogaster* data. VS developed the analytical approximation of expected transition rates. SR performed the analysis of Dobzhansky-Muller incompatibilities. RCD planned the study. RCD and RN supervised the study. JS and RCD wrote the manuscript. All authors read and edited the manuscript.

## Acknowledgments

This work was supported by the Institute of General Medical Sciences at the National Institutes (grant number R35GM128932) and an award from the Alfred P. Sloan Foundation to RBC. RN, RCD were funded within the framework of the HSE University Basic Research Program. VS was supported by grant RFBR 19-07-00515.

## Data availability statement

The source code of Ancestry_HMM-S can be downloaded from https://github.com/jesvedberg/Ancestry_HMM-S/. A user manual is also available at this location. No new data was generated for this study.

## Supplementary materials

### Supplementary figures

**Figure S1:**
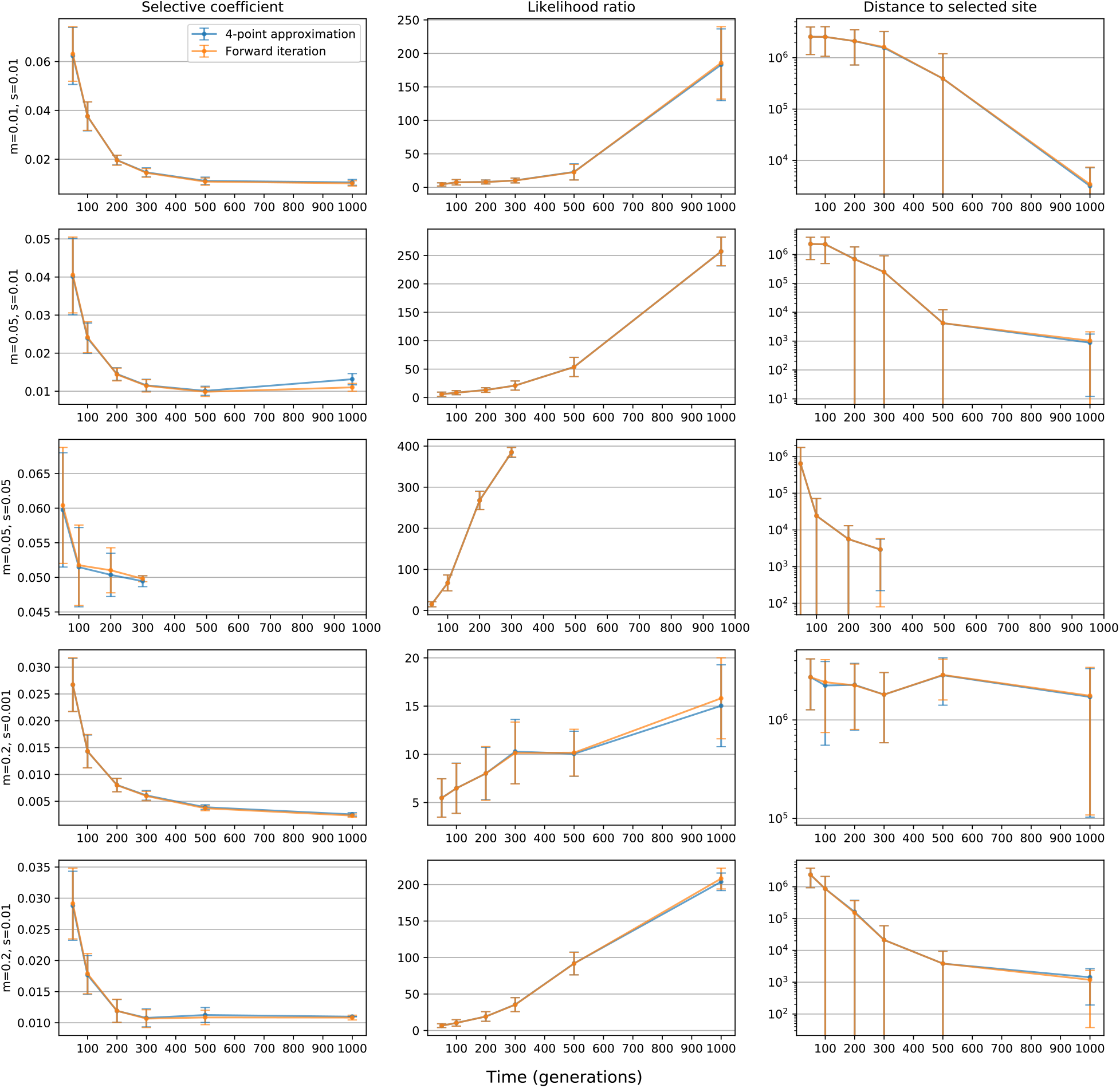
Comparison of the two different algorithms implemented in AHMM-S for calculating Markovian transition rates surrounding a selected locus. The forward iteration method will iterate through each site on a chromosome and calculate the expected transition rates between them assuming a logistic allele frequency trajectory for the selected site. In the faster 4-point analytical approximation, only the transition rates between 4 points on each side of selected sites are calculated, and the transition rates of the rest of the sites can then be interpolated. These two methods perform in close to identical fashion over all parameters that have been compared and we therefore use the 4-point approximation as our default to facilitate rapid computation.

**Figure S2:**
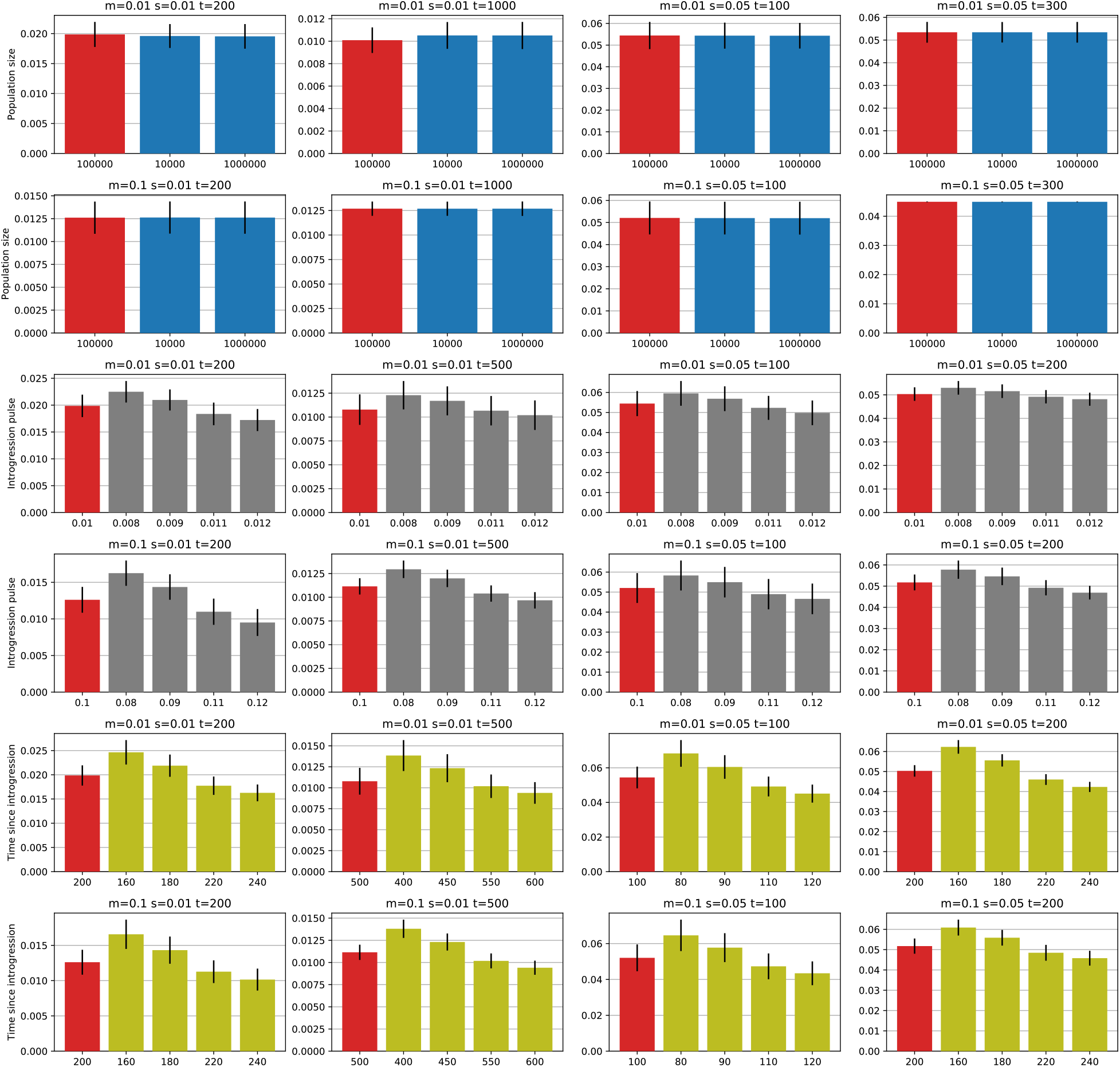
Effects of misspecifying the size of the population, the size of the introgression pulse and the time since introgression in generations on the estimated selective coefficient. The true value is shown in red. The model is robust for misspecification of population size (blue bars), where using a value an order of magnitude larger or smaller has insignificant effects. For the size (grey bars) and timing of introgression (green bars), the effects are larger, with misspecifications of ~20% generating errors of 20-40%. Overestimating these parameters leads to an underestimated value of s, and vice versa.

**Figure S3:**
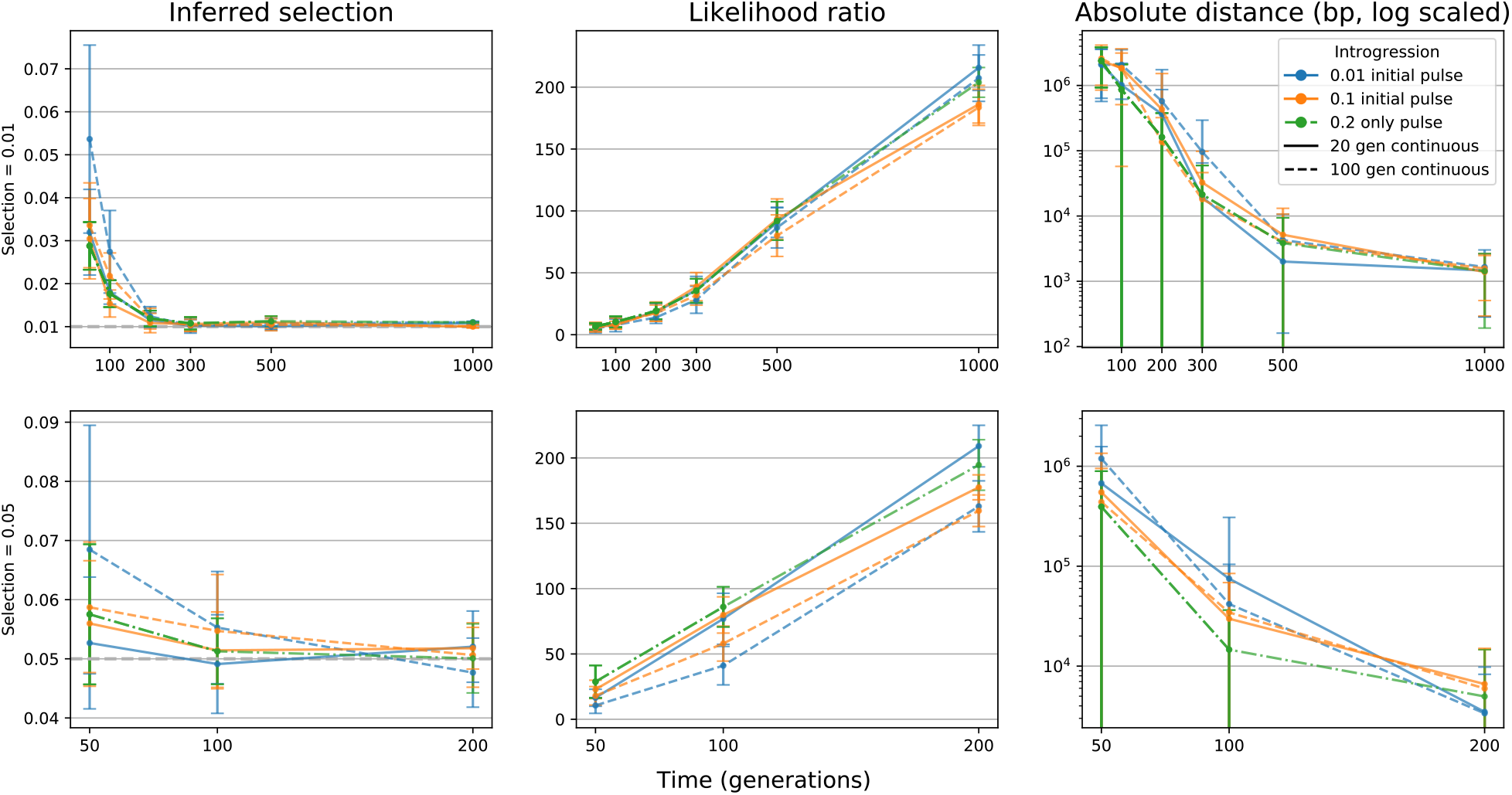
Continuous gene flow. We performed simulations where after an initial introgression pulse of 1% or 10%, gene flow continued at 1% per generation for 20 generations, or 0.2% per generation for 100 generations. The selected allele had s=0.01 or s=0.05. The final background introgression fraction was in all cases ~20%, and we compare these simulations to a single 20% pulse. The x axis shows time in generation since introgression and the y axis either the inferred value of s, the likelihood ratio or the distance between the inferred position and the real position. Continuous gene flow had little effect on both the ability to identify the correct locus and on estimating the selective coefficient. At early time points, especially when gene flow was still ongoing, estimates of *s* was higher than with a single introgressive pulse, but from 100-200 generations, the differences were minor.

**Figure S4:**
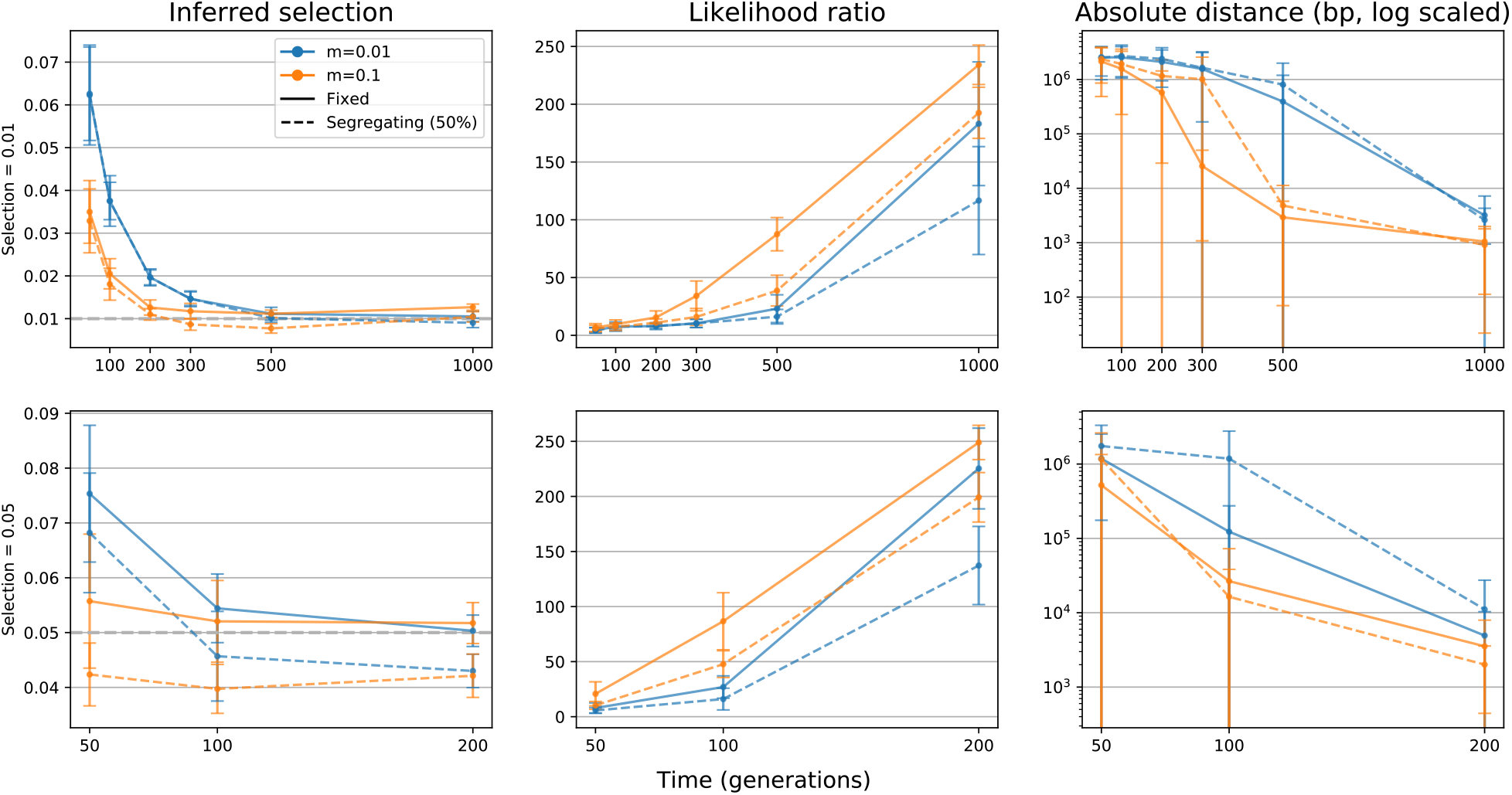
Selected site segregating in donor population. We performed simulations of an adaptive introgression scenario where the adaptive allele is segregating at 50% in the donor population. The introgression pulse in the simulations were either 1% or 10%, and s was either 0.01 or 0.05. The x axis shows time in generation since introgression and the y axis either the inferred value of s, the likelihood ratio or the distance between the inferred position and the real position. When the selected site is segregating in the donor population, AHMM-S infers the position with less accuracy at intermediate time points, but as the time since admixture increases the difference in precision compared to when the site is fixed diminishes. The selective coefficient is generally underestimated, with values up to 20% below the true value.

**Figure S5:**
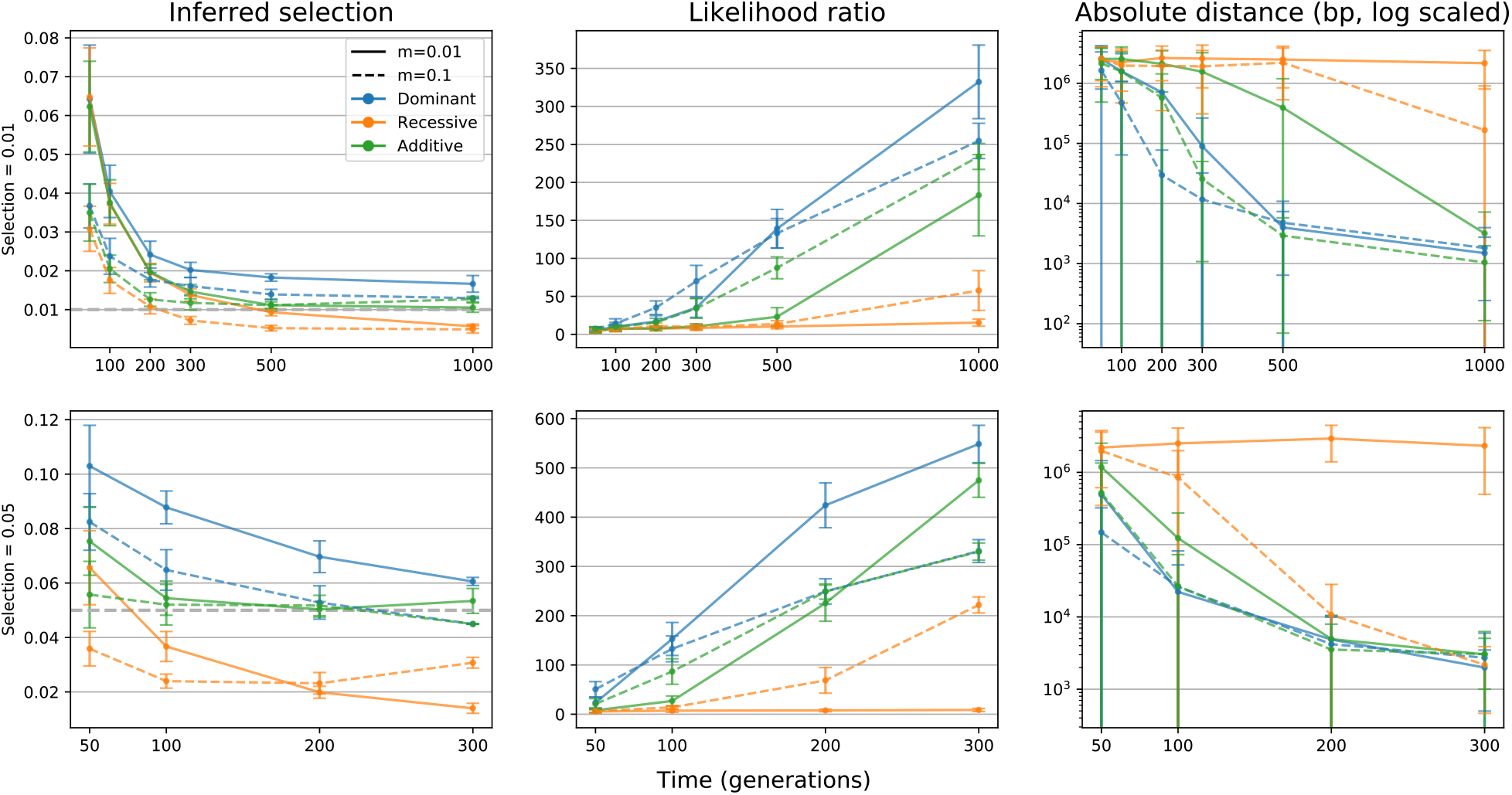
Dominant vs recessive vs additive selection. Results for simulations where the selected allele acted as a dominant, recessive or additive locus. (All other simulations use additive selection). Introgression pulse in the simulations was either set to 1% or 10% and the selection coefficient s to 0.01 or 0.05. The x axis shows time in generation since introgression and the y axis either the inferred value of s, the likelihood ratio or the distance between the inferred position and the real position. Dominant selection appears to improve the precision in identifying the correct position (probably due to increasing in frequency more quickly), but the estimated value of *s* is generally higher than the true value. An allele that acts recessively will conversely show the opposite pattern, most likely because it increases in frequency more slowly. Especially in the case where the introgressive pulse is only 1%, we are never able to identify the site with adequate precision, probably due to the introgressed allele only very rarely being found in a homozygous state.

**Figure S6:**
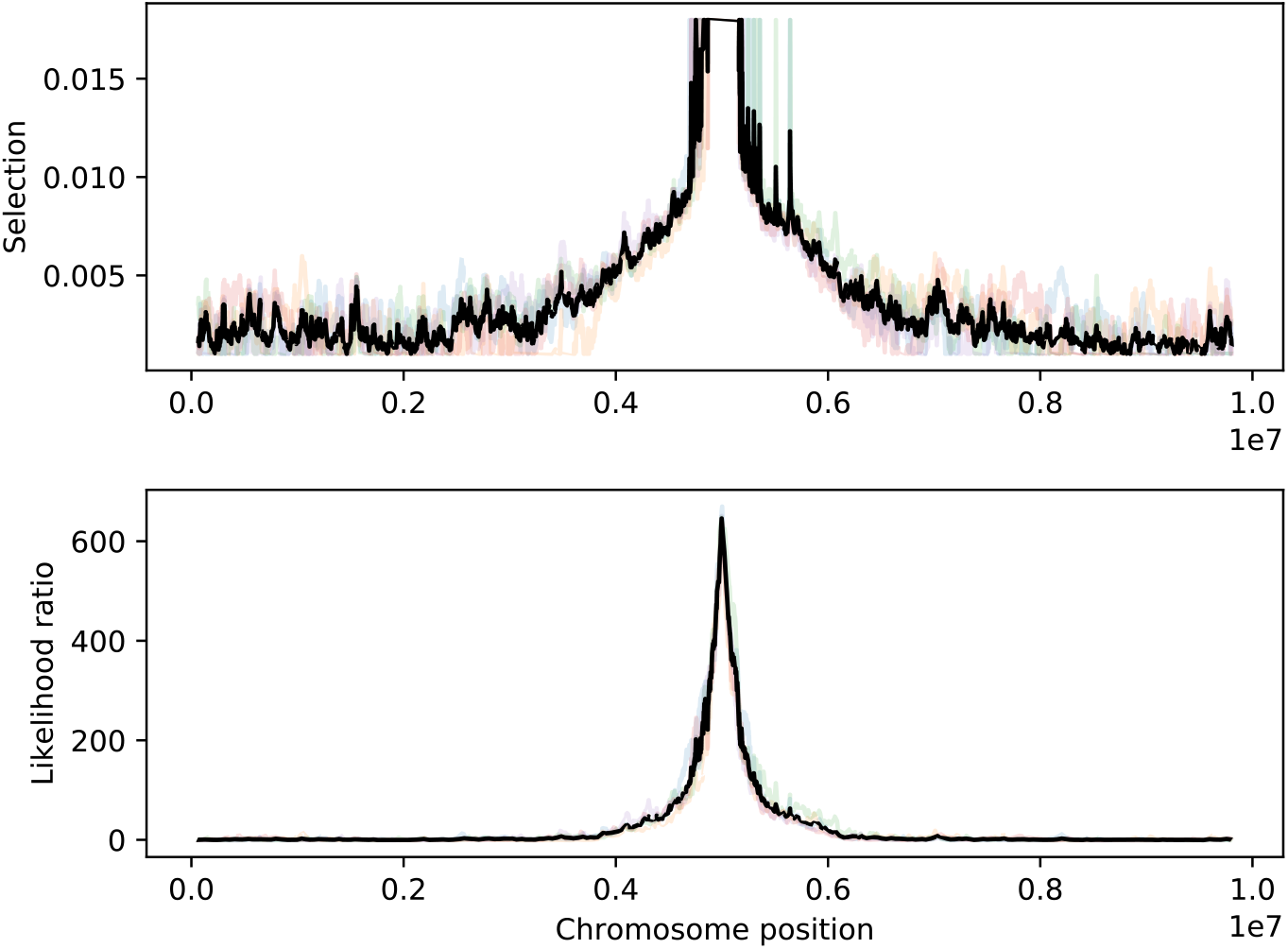
Example of AHMM-S analysing an adaptive introgression scenario where the selected site has gone to fixation. Inferred selected coefficient and likelihood ratio from five simulations (and the mean of the five simulations) of a 1% introgression of an allele with s=0.05, sampled after 1000 generations. The likelihood ratio peak identifies the correct position with high accuracy, but AHMM-S will only report a peak selection coefficient of ~0.017, as that is the value of s that causes a site to go to fixation at 1000 generations.

**Figure S7:**
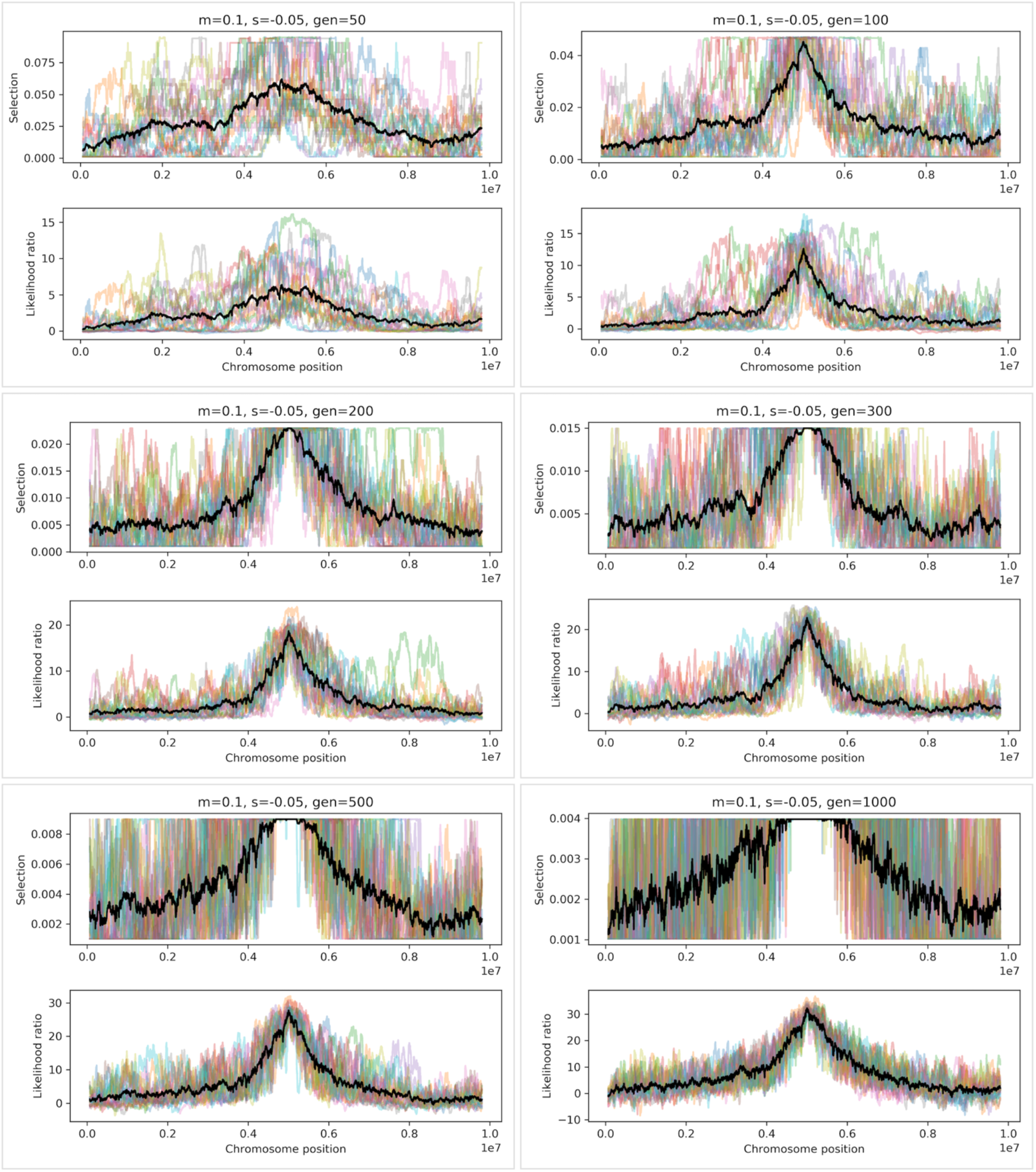
Negative selection. Example of a simulation of an introgressed site experiencing negative selection, with an introgressive fraction of 0.1 and a selective coefficient of 0.05. We ran 20 simulations and sampled them from 50 to 1000 generations. We analyzed the simulations with AHMM-S after reverting the genotype identity, so the negative selection of the introgressed site shows up as positive selection of the receiving genotype. AHMM-S can identify the location of the selected site, but as the frequency of the site over time approaches zero, the precision of the estimated selective coefficient is lost.

**Figure S8:**
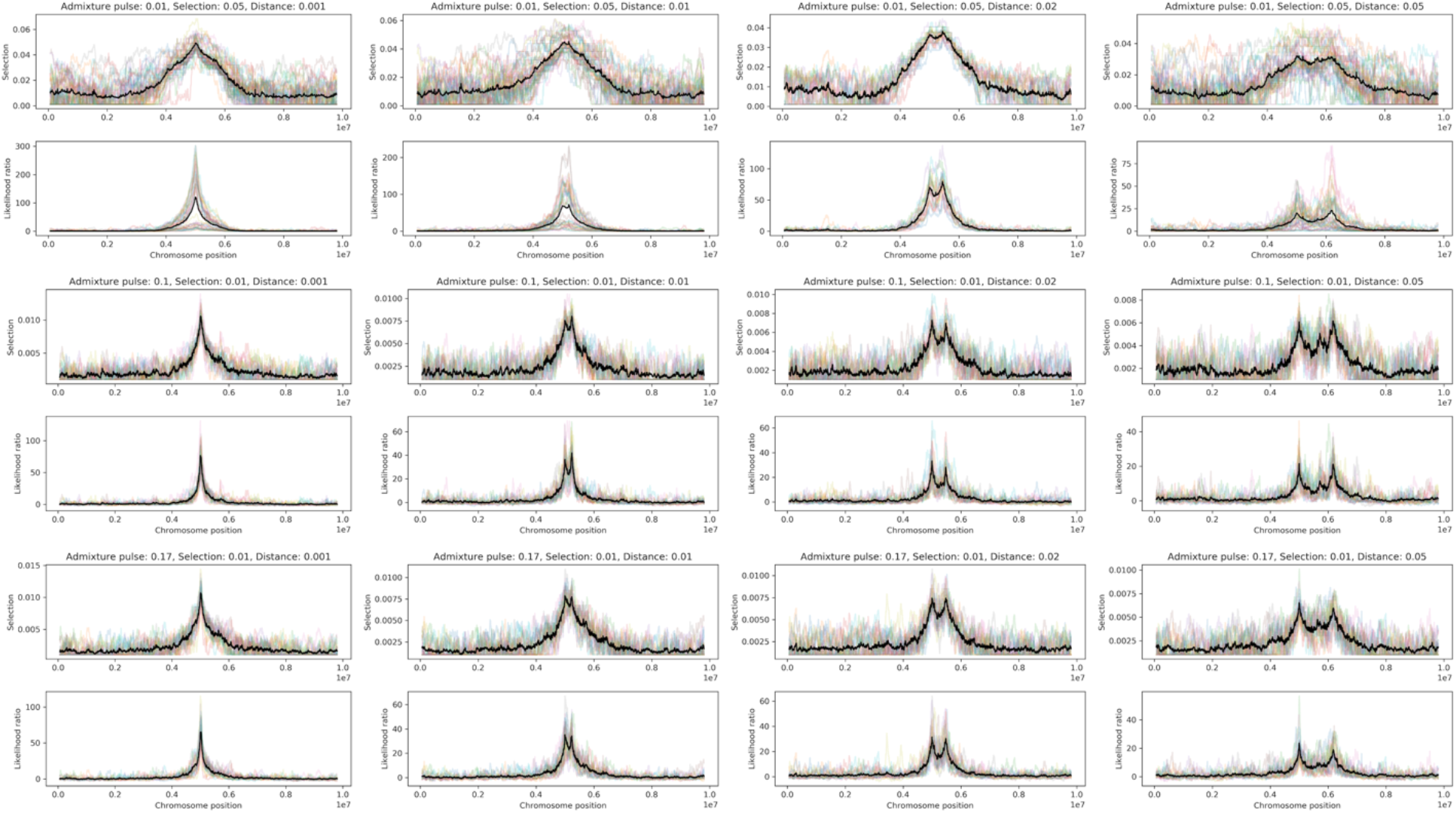
Peak separation for closely linked selected sites. Three different introgression scenarios were simulated (A: m=0.1, s=0.01 t=500; B: m=0.01, s=0.05, t=200; C: m=0.17, s=0.01, t=430) in 20 replicates with two sites of equal selective strength (s/2) located at increasing distances (0.1, 1, 2 and 5 cM) from each other. Two distinct peaks are visible in the cases of 1, 2 and 5 cM for all three scenarios.

**Figure S9:**
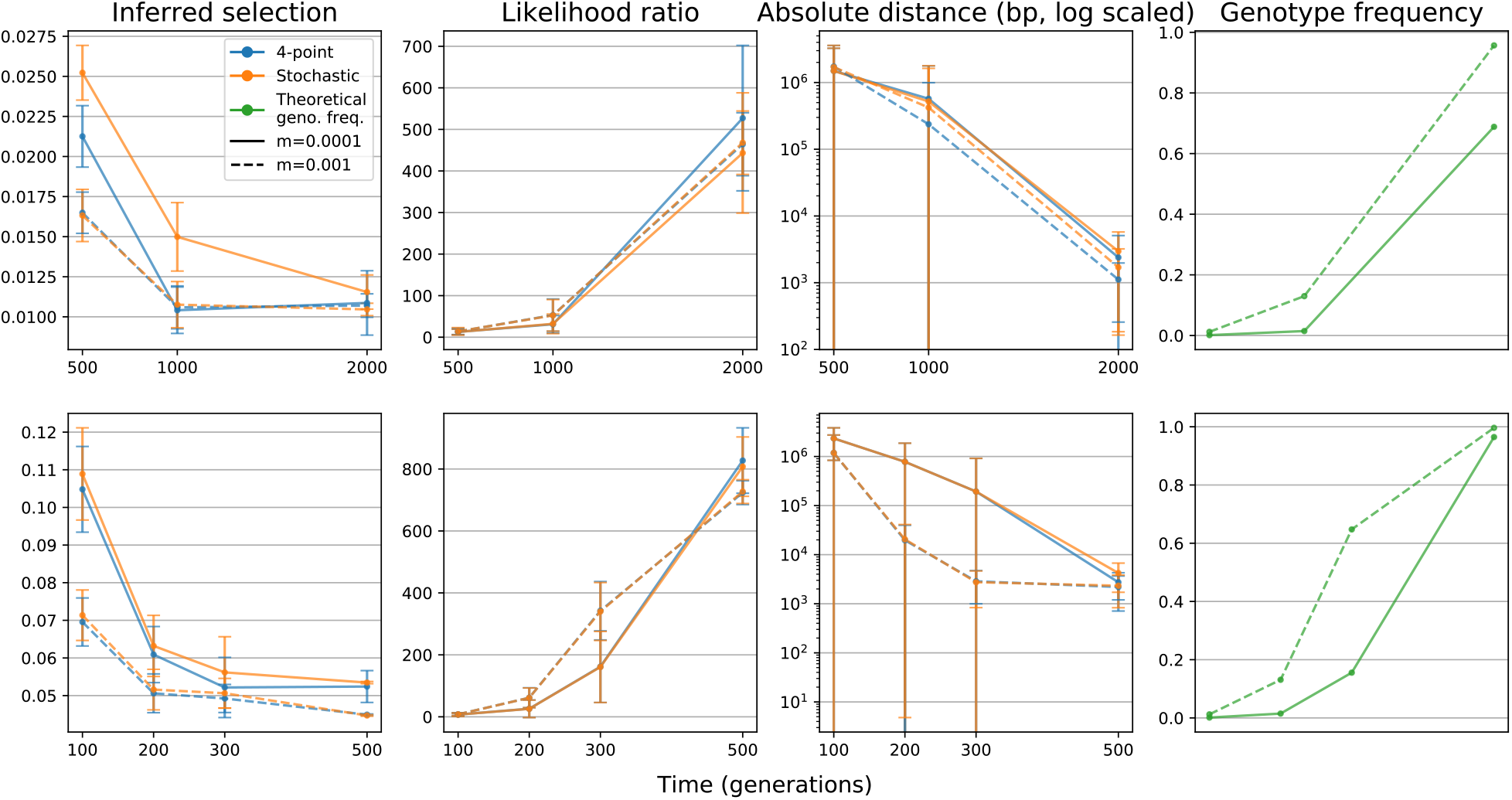
Small introgression fractions. Results for simulations with a low introgression fraction. We compared the performance of the 4-point approximative algorithm for calculating the expected increase in frequency of an adaptively introgressed allele, with a stochastic algorithm based on averaging multiple forward simulations. Small introgression fractions is expected to violate the model used for the 4-point approximation, but it performs equally well as the more precise (but significantly slower) stochastic method.

**Figure S10:**
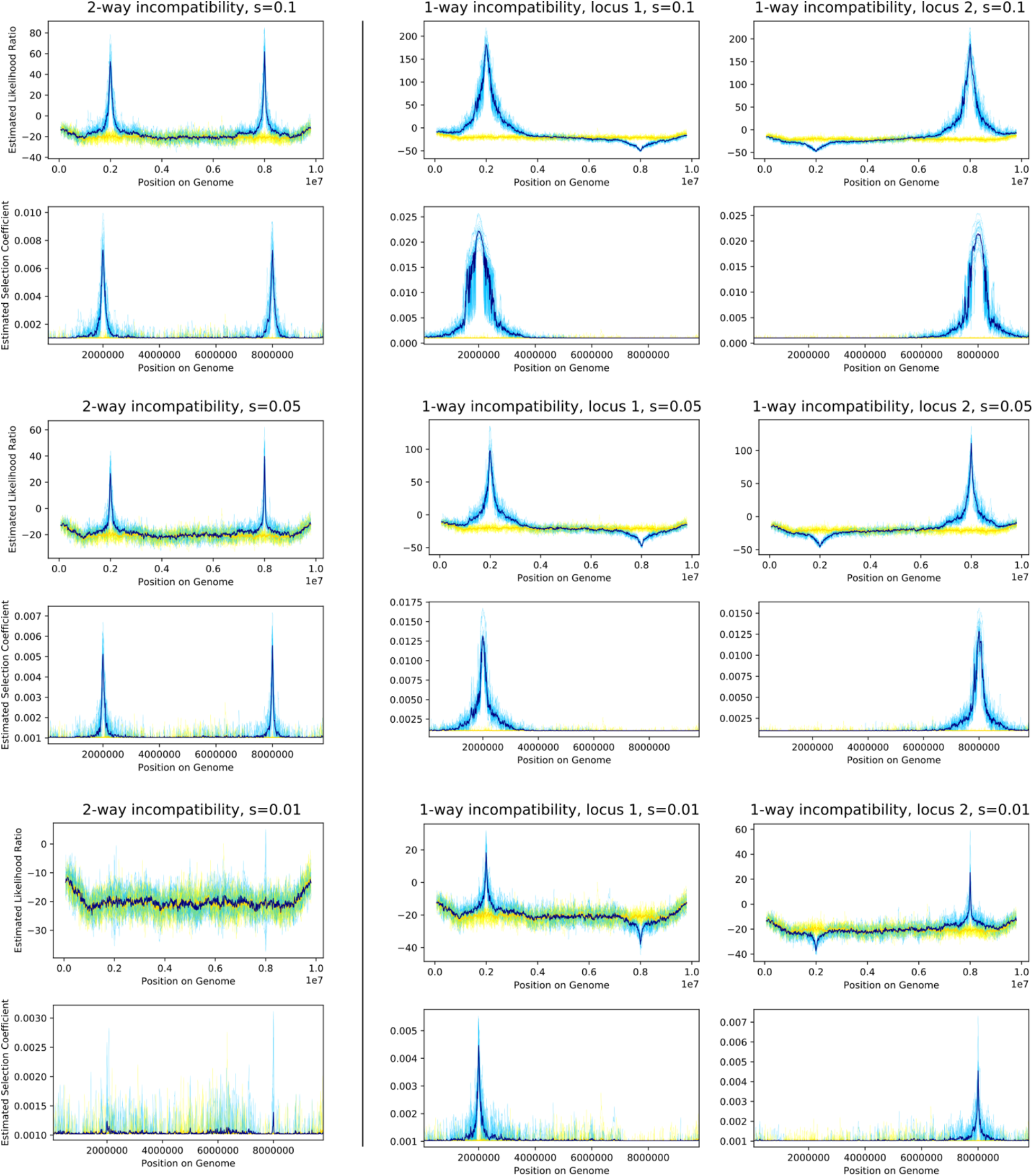
Dobzhansky-Muller Incompatibilities (DMIs) identified using AHMM-S. Two scenarios were investigated for three different values of s (−0.1, −0.05 and −0.01) with an introgressive pulse of 0.5. In both scenarios, two loci (A and B), located at 2 Mbp and 8 Mbp, have an incompatibility interaction between alleles from the two different populations. In scenario 1 (column 1), any interaction between different alleles is selected against (A0B1 and A1B0), in scenario 2 (column 2 and 3) only one combination (A0B1) is selected against. The interacting loci could be identified in all cases except scenario 1, s=0.01. The results for scenario 2 are split into two columns as the direction of selection is different for each locus and AHMM-S must identify each locus separately in different analyses. Blue lines show simulations with DMI and yellow lines negative controls without any active selection. Darker lines show mean values and lighter lines the results for each simulation. Negative likelihood ratios can appear when the likeliest inferred value of s is less likely than the neutral case. This can be explained by the search space for s starting at 0.001.

**Figure S11:**
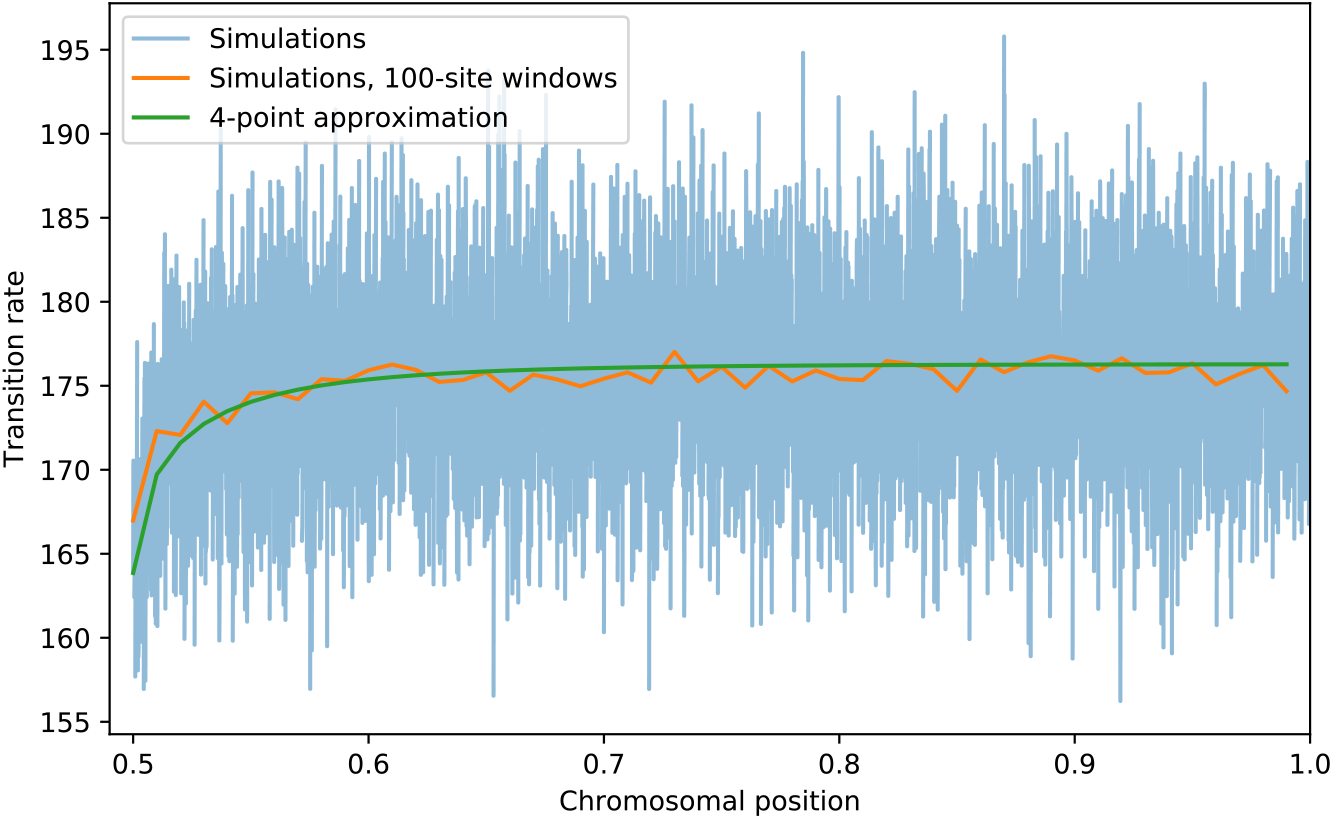
Simulated vs expected transition rates. Transition rates for 100,000 simulations are plotted at each site (blue), or averaged in 100-site windows (orange), and compared to transition rates estimated with the 4-point approximation method (green). The selected site is located at position 0.5 and the transition rates are plotted going away from the selected site to the end of the chromosome at position 1.

### Supplementary tables

**Table S1:** GO categories including more than 10 candidate genes

| GO Term | Number of genes | Q-value (FDR corrected) | Description | Genes Detected |
| --- | --- | --- | --- | --- |
| GO:0097159 | 21 | 0.094285 | organic cyclic compound binding | tRNA:Arg-ACG-1-4, osa, Cyp6a22, Cyp6w1, tRNA:Lys-CTT-1-5, RIOK2, tRNA:Arg-ACG-1-3, tRNA:Arg-ACG-1-5, Cyp6a17, trx, Cyp6g1, SmF, scaRNA:MeU2-C41, dsx, Pcf11, EndoG, GlnRS, CG10445, CG2678, lds, cic |
| GO:1901363 | 21 | 0.094285 | heterocyclic compound binding | tRNA:Arg-ACG-1-4, osa, Cyp6a22, Cyp6w1, tRNA:Lys-CTT-1-5, RIOK2, tRNA:Arg-ACG-1-3, tRNA:Arg-ACG-1-5, Cyp6a17, trx, Cyp6g1, SmF, scaRNA:MeU2-C41, and dsx, Pcf11, EndoG, GlnRS, CG10445, CG2678, lds, cic |
| GO:0003676 | 13 | 0.4480193103 | nucleic acid binding | tRNA:Arg-ACG-1-4, osa, tRNA:Lys-CTT-1-5, tRNA:Arg-ACG-1-3, tRNA:Arg-ACG-1-5, trx, SmF, scaRNA:MeU2-C41, dsx, Pcf11, EndoG, CG2678, cic |

## References

Aldridge WN. 1950. Some properties of specific cholinesterase with particular reference to the mechanism of inhibition by diethyl p-nitrophenyl thiophosphate (E 605) and analogues. Biochem. J. 46:451–460.

Alexander DH, Novembre J, Lange K. 2009. Fast model-based estimation of ancestry in unrelated individuals. Genome Res. 19:1655–1664.

Aminetzach YT, Macpherson JM, Petrov DA. 2005. Pesticide Resistance via Transposition-Mediated Adaptive Gene Truncation in Drosophila. Science 309:764–767.

Baran Y, Pasaniuc B, Sankararaman S, Torgerson DG, Gignoux C, Eng C, Rodriguez-Cintron W, Chapela R, Ford JG, Avila PC, et al. 2012. Fast and accurate inference of local ancestry in Latino populations. Bioinformatics 28:1359–1367.

Battlay P, Leblanc PB, Green L, Garud NR, Schmidt JM, Fournier-Level A, Robin C. 2018. Structural Variants and Selective Sweep Foci Contribute to Insecticide Resistance in the Drosophila Genetic Reference Panel. G3 Genes Genomes Genet. 8:3489–3497.

Begun DJ, Aquadro CF. 1992. Levels of naturally occurring DNA polymorphism correlate with recombination rates in D. melanogaster. Nature 356:519–520.

Bergland AO, Tobler R, González J, Schmidt P, Petrov D. 2016. Secondary contact and local adaptation contribute to genome-wide patterns of clinal variation in Drosophila melanogaster. Mol. Ecol. 25:1157–1174.

Biscoe ML, Kramer RA, Mutero CM. 2005. Current Policy and Status of DDT Use for Malaria Control in Ethiopia, Uganda, Kenya and South Africa. International Water Management Institute Available from: https://dukespace.lib.duke.edu/dspace/handle/10161/7481

Chen GK, Marjoram P, Wall JD. 2009. Fast and flexible simulation of DNA sequence data. Genome Res. 19:136–142.

Chowdhuri DK, Nazir A, Saxena DK. 2001. Effect of three chlorinated pesticides on hsromega stress gene in transgenic Drosophila melanogaster. J. Biochem. Mol. Toxicol. 15:173–186.

Comeron JM, Ratnappan R, Bailin S. 2012. The Many Landscapes of Recombination in Drosophila melanogaster. PLOS Genet 8:e1002905.

Corbett-Detig R, Jones M. 2016. SELAM: simulation of epistasis and local adaptation during admixture with mate choice. Bioinformatics 32:3035–3037.

Corbett-Detig R, Nielsen R. 2017. A Hidden Markov Model Approach for Simultaneously Estimating Local Ancestry and Admixture Time Using Next Generation Sequence Data in Samples of Arbitrary Ploidy. PLoS Genet. [Internet] 13. Available from: https://www.ncbi.nlm.nih.gov/pmc/articles/PMC5242547/

Corbett-Detig RB, Hartl DL. 2012. Population Genomics of Inversion Polymorphisms in Drosophila melanogaster. PLOS Genet. 8:e1003056.

Corbett-Detig RB, Zhou J, Clark AG, Hartl DL, Ayroles JF. 2013. Genetic incompatibilities are widespread within species. Nature 504:135–137.

Coyne JA, Orr HA. 2004. Speciation. Sinauer

Daborn PJ, Yen JL, Bogwitz MR, Goff GL, Feil E, Jeffers S, Tijet N, Perry T, Heckel D, Batterham P, et al. 2002. A Single P450 Allele Associated with Insecticide Resistance in Drosophila. Science 297:2253–2256.

Falush D, Stephens M, Pritchard JK. 2003. Inference of population structure using multilocus genotype data: linked loci and correlated allele frequencies. Genetics 164:1567–1587.

Fraïsse C, Roux C, Welch JJ, Bierne N. 2014. Gene-Flow in a Mosaic Hybrid Zone: Is Local Introgression Adaptive? Genetics 197:939–951.

Garud NR, Messer PW, Buzbas EO, Petrov DA. 2015. Recent Selective Sweeps in North American Drosophila melanogaster Show Signatures of Soft Sweeps. PLoS Genet. [Internet] 11. Available from: https://www.ncbi.nlm.nih.gov/pmc/articles/PMC4338236/

Gower G, Picazo PI, Fumagalli M, Racimo F. 2020. Detecting adaptive introgression in human evolution using convolutional neural networks. bioRxiv:2020.09.18.301069.

Gravel S. 2012. Population Genetics Models of Local Ancestry. Genetics 191:607–619.

Harris K, Nielsen R. 2016. The Genetic Cost of Neanderthal Introgression. Genetics 203:881–891.

Hedrick PW. 2013. Adaptive introgression in animals: examples and comparison to new mutation and standing variation as sources of adaptive variation. Mol. Ecol. 22:4606–4618.

Huerta-Sánchez E, Jin X, Asan, Bianba Z, Peter BM, Vinckenbosch N, Liang Y, Yi X, He M, Somel M, et al. 2014. Altitude adaptation in Tibetans caused by introgression of Denisovan-like DNA. Nature 512:194–197.

Jeong C, Alkorta-Aranburu G, Basnyat B, Neupane M, Witonsky DB, Pritchard JK, Beall CM, Di Rienzo A. 2014. Admixture facilitates genetic adaptations to high altitude in Tibet. Nat. Commun. 5:3281.

Kao JY, Lymer S, Hwang SH, Sung A, Nuzhdin SV. 2015. Postmating reproductive barriers contribute to the incipient sexual isolation of the United States and Caribbean Drosophila melanogaster. Ecol. Evol. 5:3171–3182.

Kaplan NL, Hudson RR, Langley CH. 1989. The ‘‘hitchhiking Effect’’ Revisited. Genetics 123:887–899.

Karasov T, Messer PW, Petrov DA. 2010. Evidence that Adaptation in Drosophila Is Not Limited by Mutation at Single Sites. PLOS Genet. 6:e1000924.

Kim BY, Huber CD, Lohmueller KE. 2018. Deleterious variation shapes the genomic landscape of introgression. PLOS Genet. 14:e1007741.

Kofler R, Schlötterer C. 2012. Gowinda: unbiased analysis of gene set enrichment for genomewide association studies. Bioinformatics 28:2084–2085.

Kolaczkowski B, Kern AD, Holloway AK, Begun DJ. 2011. Genomic Differentiation Between Temperate and Tropical Australian Populations of Drosophila melanogaster. Genetics 187:245–260.

Lack JB, Cardeno CM, Crepeau MW, Taylor W, Corbett-Detig RB, Stevens KA, Langley CH, Pool JE. 2015. The Drosophila Genome Nexus: A Population Genomic Resource of 623 Drosophila melanogaster Genomes, Including 197 from a Single Ancestral Range Population. Genetics 199:1229–1241.

Lack JB, Lange JD, Tang AD, Corbett-Detig RB, Pool JE. 2016. A Thousand Fly Genomes: An Expanded Drosophila Genome Nexus. Mol. Biol. Evol. 33:3308–3313.

Langley CH, Stevens K, Cardeno C, Lee YCG, Schrider DR, Pool JE, Langley SA, Suarez C, Corbett-Detig RB, Kolaczkowski B, et al. 2012. Genomic Variation in Natural Populations of Drosophila melanogaster. Genetics 192:533–598.

Liang M, Nielsen R. 2014. The Lengths of Admixture Tracts. Genetics 197:953–967.

Loh P-R, Lipson M, Patterson N, Moorjani P, Pickrell JK, Reich D, Berger B. 2013. Inferring Admixture Histories of Human Populations Using Linkage Disequilibrium. Genetics 193:1233–1254.

Lohmueller KE, Bustamante CD, Clark AG. 2011. Detecting Directional Selection in the Presence of Recent Admixture in African-Americans. Genetics 187:823–835.

Magwire MM, Bayer F, Webster CL, Cao C, Jiggins FM. 2011. Successive Increases in the Resistance of Drosophila to Viral Infection through a Transposon Insertion Followed by a Duplication. PLOS Genet. 7:e1002337.

Maples BK, Gravel S, Kenny EE, Bustamante CD. 2013. RFMix: A Discriminative Modeling Approach for Rapid and Robust Local-Ancestry Inference. Am. J. Hum. Genet. 93:278–288.

Marjoram P, Wall JD. 2006. Fast “coalescent” simulation. BMC Genet. 7:16.

Medina P, Thornlow B, Nielsen R, Corbett-Detig R. 2018. Estimating the Timing of Multiple Admixture Pulses During Local Ancestry Inference. Genetics 210:1089–1107.

Meiklejohn CD, Landeen EL, Gordon KE, Rzatkiewicz T, Kingan SB, Geneva AJ, Vedanayagam JP, Muirhead CA, Garrigan D, Stern DL, et al. 2018. Gene flow mediates the role of sex chromosome meiotic drive during complex speciation. Przeworski M, Tautz D, editors. eLife 7:e35468.

Menozzi P, Shi MA, Lougarre A, Tang ZH, Fournier D. 2004. Mutations of acetylcholinesterase which confer insecticide resistance in Drosophila melanogaster populations. BMC Evol. Biol. 4:4.

Norris LC, Main BJ, Lee Y, Collier TC, Fofana A, Cornel AJ, Lanzaro GC. 2015. Adaptive introgression in an African malaria mosquito coincident with the increased usage of insecticide-treated bed nets. Proc. Natl. Acad. Sci. [Internet]. Available from: https://www.pnas.org/content/early/2015/01/02/1418892112

Pavlidis P, Alachiotis N. 2017. A survey of methods and tools to detect recent and strong positive selection. J. Biol. Res.-Thessalon. 24:7.

Pool JE. 2015. The Mosaic Ancestry of the Drosophila Genetic Reference Panel and the D. melanogaster Reference Genome Reveals a Network of Epistatic Fitness Interactions. Mol. Biol. Evol. 32:3236–3251.

Pool JE, Corbett-Detig RB, Sugino RP, Stevens KA, Cardeno CM, Crepeau MW, Duchen P, Emerson JJ, Saelao P, Begun DJ, et al. 2012. Population Genomics of Sub-Saharan Drosophila melanogaster: African Diversity and Non-African Admixture. PLoS Genet. [Internet] 8. Available from: https://www.ncbi.nlm.nih.gov/pmc/articles/PMC3527209/

Pool JE, Nielsen R. 2009. Inference of Historical Changes in Migration Rate From the Lengths of Migrant Tracts. Genetics 181:711–719.

Powell DL, García-Olazábal M, Keegan M, Reilly P, Du K, Díaz-Loyo AP, Banerjee S, Blakkan D, Reich D, Andolfatto P, et al. 2020. Natural hybridization reveals incompatible alleles that cause melanoma in swordtail fish. Science 368:731–736.

Pritchard JK, Stephens M, Donnelly P. 2000. Inference of Population Structure Using Multilocus Genotype Data. Genetics 155:945–959.

Quinn LP, B, Vos J de, Fernandes-Whaley M, Roos C, Bouwman H, Kylin H, Pieters R, Berg J van den. 2011. Pesticide Use in South Africa: One of the Largest Importers of Pesticides in Africa. Pestic. Mod. World - Pestic. Use Manag. [Internet]. Available from: https://www.intechopen.com/books/pesticides-in-the-modern-world-pesticides-use-and-management/pesticide-use-in-south-africa-one-of-the-largest-importers-of-pesticides-in-africa

Racimo F, Sankararaman S, Nielsen R, Huerta-Sánchez E. 2015. Evidence for archaic adaptive introgression in humans. Nat. Rev. Genet. 16:359–371.

Reinhardt JA, Kolaczkowski B, Jones CD, Begun DJ, Kern AD. 2014. Parallel Geographic Variation in Drosophila melanogaster. Genetics 197:361–373.

Sankararaman S, Mallick S, Dannemann M, Prüfer K, Kelso J, Pääbo S, Patterson N, Reich D. 2014. The genomic landscape of Neanderthal ancestry in present-day humans. Nature 507:354–357.

Sankararaman S, Sridhar S, Kimmel G, Halperin E. 2008. Estimating Local Ancestry in Admixed Populations. Am. J. Hum. Genet. 82:290–303.

Schmidt JM, Battlay P, Gledhill-Smith RS, Good RT, Lumb C, Fournier-Level A, Robin C. 2017. Insights into DDT Resistance from the Drosophila melanogaster Genetic Reference Panel. Genetics 207:1181–1193.

Schumer M, Cui R, Powell DL, Dresner R, Rosenthal GG, Andolfatto P. 2014. High-resolution mapping reveals hundreds of genetic incompatibilities in hybridizing fish species. McVean G, editor. eLife 3:e02535.

Schumer M, Powell DL, Corbett-Detig R. 2020. Versatile simulations of admixture and accurate local ancestry inference with mixnmatch and ancestryinfer. Mol. Ecol. Resour. [Internet] n/a. Available from: https://onlinelibrary.wiley.com/doi/abs/10.1111/1755-0998.13175

Setter D, Mousset S, Cheng X, Nielsen R, DeGiorgio M, Hermisson J. 2020. VolcanoFinder: Genomic scans for adaptive introgression. PLOS Genet. 16:e1008867.

Shchur V, Svedberg J, Medina P, Corbett-Detig R, Nielsen R. 2020. On the Distribution of Tract Lengths During Adaptive Introgression. G3 Genes Genomes Genet. 10:3663–3673.

Song Y, Endepols S, Klemann N, Richter D, Matuschka F-R, Shih C-H, Nachman MW, Kohn MH. 2011. Adaptive introgression of anticoagulant rodent poison resistance by hybridization between Old World mice. Curr. Biol. CB 21:1296–1301.

Suarez-Gonzalez, Adriana, Lexer, Christian, Cronk, Quentin C. B. 2018. Adaptive introgression: a plant perspective. Biol. Lett. 14:20170688.

The Heliconius Genome Consortium. 2012. Butterfly genome reveals promiscuous exchange of mimicry adaptations among species. Nature 487:94–98.

Thornton K, Andolfatto P. 2006. Approximate Bayesian Inference Reveals Evidence for a Recent, Severe Bottleneck in a Netherlands Population of Drosophila melanogaster. Genetics 172:1607–1619.

Vernot B, Akey JM. 2014. Resurrecting Surviving Neandertal Lineages from Modern Human Genomes. Science 343:1017–1021.

